# Database Permeating through Time, Space, and Medicine: A sequence, structure, and clinical compilation and comparison of transmembrane amino acids in VGL ion channels

**DOI:** 10.1101/2025.01.10.630794

**Authors:** Frank Yeh, Arya Patel, Matthew Mun, Youjin Oh, Richard W. Aldrich

## Abstract

Ion channels of the VGL superfamily are extremely diverse in their physiological roles – critical in all excitable cells. Mutations in these channels can cause clinical deficits and symptomologies. With recent advances in sequencing and structural data in the past 30 years, there is a plethora of bioinformatic information that experimentalists can utilize to draw hypotheses about structure and function. However, there is no cohesive database that has compiled this data and presents it in an easily accessible and avoids the confounds of close evolutionary lineages. Here, we seek to fill this gap in resources by compiling and comparing groups of sequences and structures across many channels of the VGL superfamily. Additionally, we corroborate the well-conserved amino acid positions with previous experimental work as well as put forth some understudied but well-conserved amino acid positions that might be fundamental to the certain generalizable mechanisms present in the VGL superfamily channels. To further lend evidence to understudied positions being critical to certain generalizable mechanisms, we also compiled clinical mutation data at these positions across many channels to show the likely functional relevance of these positions. Finally, we will make all of the alignments and resources we generated publicly available at https://github.com/Frank-Yeh-95/VGLDatabaseCompilation for ease of hypothesis generation.

## Introduction

Channels of the voltage-gated-like superfamily (VGL) are diverse in their physiological roles. To accommodate for their varied physiological niches, these channels can be categorized in different ways such as preferred stimulus, ion selectivity, and domain-swapping architecture. Advancements in sequencing and structural determination technologies have drastically increased our understanding of these channels, lending to a rich amount of bioinformatics to be compiled and analyzed. Historically, bioinformatics data has been used to draw hypotheses about the important functional contributions of specific highly conserved amino acids. However, there are few cohesive databases that can be used to investigate broadly across the VGL superfamily. Existing work has focused on extensively cataloging the VGL ion channel superfamily and giving a detailed description of the physiological role of each ion channel (Yu et al., 2005; Yu and Catterall, 2004; Zheng and Trudeau, 2015; Nilius and Owsianik, 2011; Hille, 2001; Rudy et al., 2009; Ranjan et al., 2011), but does not individually describe each of the highly conserved amino acids. More recently, sequence and structural alignments have been performed on TRP channels, a subfamily of the VGL superfamily, to generate hypotheses on generalizable mechanisms of TRP channel structure-function and on specific TRP channel amino acid differences (Cabezas-Bratesco et al., 2022; Huffer et al., 2020).

To address the gap in data compilation in other subfamilies of the VGL superfamily, we performed a sequence and structural alignment of all non-TRP, homotetrameric six transmembrane- segmented (6TMS) or twenty-four transmembrane-segmented (24TMS) channels in this superfamily. In this study, we will mention well-studied and well-conserved amino acid positions, corroborating sequence, structure, clinical and functional data. We will highlight well-conserved, but less studied amino acid positions that could be key to understanding broadly shared functional or structural mechanisms, warranting further investigations. Overall, we intend to build a database from which hypotheses of broad scope or of specific VGL channel members can be drawn.

Although the channels of the VGL superfamily are diverse in their physiological roles, we chose to focus on the large population of these channels that have a core structural similarity - these channels all share six TMS as their (pseudo)monomeric representation. These six transmembrane segments can further be divided into two structural domains - the voltage-sensing domain (VSD) and the pore-gate domain (PGD). Notably, the VSD consists of transmembrane segments 1-4 (S1-4) whereas the PGD comprises transmembrane segments 5-pore-helix-6 (S5-6). The VSD is broadly responsible for voltage-sensing in voltage-gated channels, while the PGD is home to the ion permeation pathway and the gate. Given this core structural similarity and the high variability in length and amino acid identity of the linker regions between transmembrane segments, we will only investigate the sequence and structural alignments of the TMS.

The channels that fit this the 6TMS-like, (pseudo)monomeric structure (Fig. 1) are the Shaker family of voltage-gated potassium channels (Kv1-4; 6TMS), KCNQ voltage-gated potassium channels (Kv7; 6TMS), CNBD family of channels (Kv10-12, HCN, CNG; 6TMS), Slo family of potassium channels (BK, Slo2, Slo3; 6TMS), calcium-activated small-conductance potassium channels (SK, IK; 6TMS), voltage-gated calcium channels (Cav1-3; 24TMS), voltage-gated sodium channels (Nav1; 24TMS), and non-selective sodium leak channels (NALCN; 24TMS) (Yu et al., 2005; Yu and Catterall, 2004; Monteil et al., 2024; Ranjan et al., 2011; Zheng and Trudeau, 2015; Hille, 2001). These channels cover a wide breadth of physiological roles in excitable cells. Potassium channels set the resting membrane potential, repolarize the membrane after an action potential, and provide local negative feedback to voltage-sensitive calcium-influx sources (Hodgkin and Huxley, 1952a; Gómez et al., 2021; Zhang et al., 2018; Parks et al., 2017). While most voltage-gated channels gate to depolarizing potentials, HCN channels gate to hyperpolarizing potentials contributing to resetting the membrane potential after hyperpolarizations and contribute to the oscillatory properties of certain excitable cells (Mishra and Narayanan, 2025; Robinson and Siegelbaum, 2003; DiFrancesco, 2010). Voltage-gated sodium and calcium channels are responsible for the depolarization phase of the action potential (Catterall, 2011; Santana et al., 2010; Golding et al., 1999; Hodgkin and Huxley, 1952b; Wang et al., 2017) and, for calcium, the initiation of the synaptic transmission pathway as well as the control for muscle contraction (Dolphin and Lee, 2020; Pelizzari et al., 2024; Pereira da Silva et al., 2022). The non-selective sodium leak channels regulate the resting membrane potential through the sodium leak currents (Monteil et al., 2024; Lu et al., 2007). Critically, mutations in any of these channels can lead to severe developmental, mental, sensory perception, motor, and cardiac abnormalities (Suppl. Tables 3-10 and Files 9-16).

**Figure 1:**
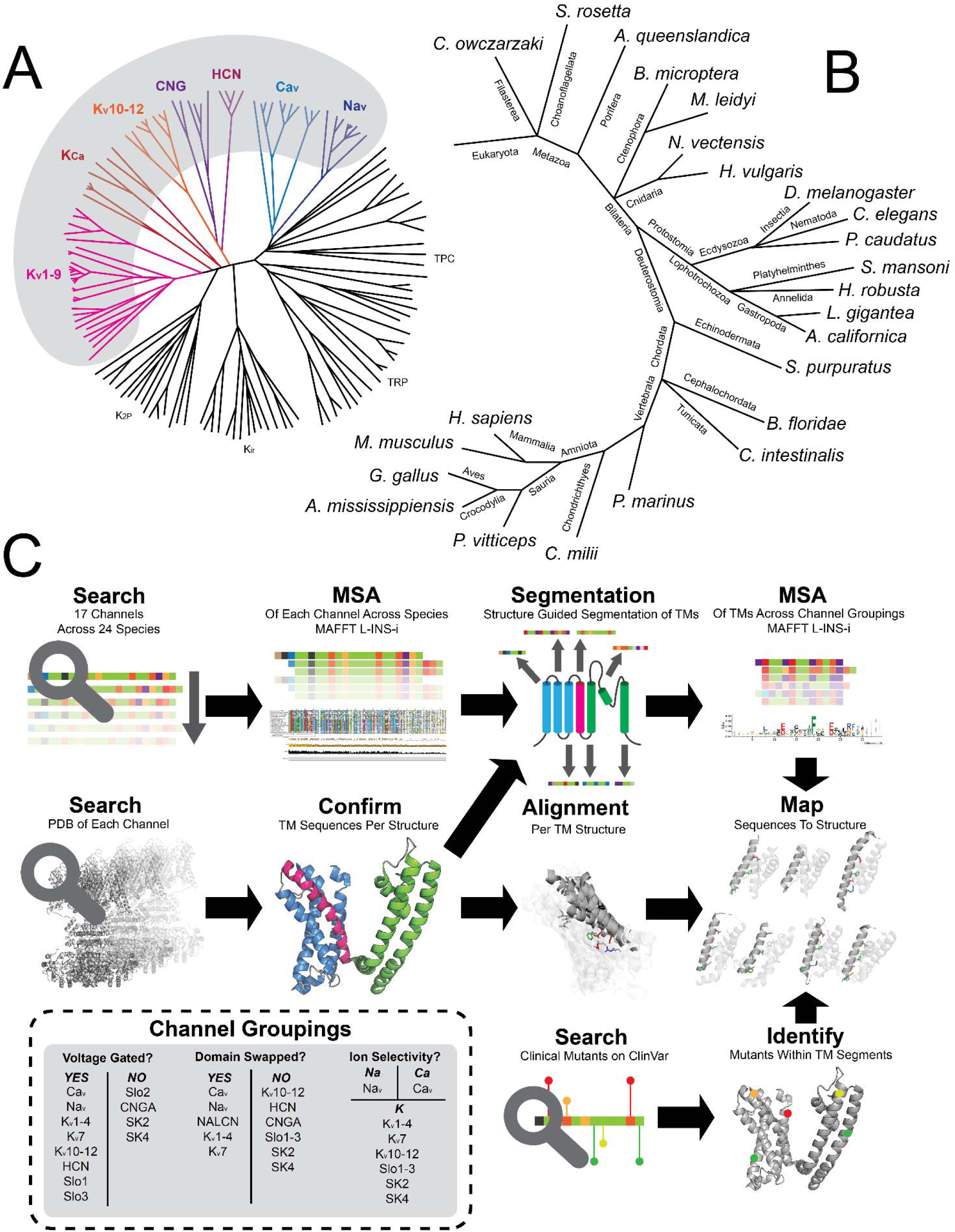
Workflow and Development of the database: **A.** is the VGL family tree as from (Yu and Catterall, 2004; Yu et al., 2005). Here, we highlight the channels that we will be including in this database compilation. In particular, we will include: Kv1-4, 7, 10-12; KCa1-5; HCN; CNGA; Cav1-3; Nav1; and NALCN channels (Yu and Catterall, 2004; Yu et al., 2005). **B.** shows the species genomes that were queried for all of the channels denoted in **A.** These species are meant to cover a wide swath of organisms across the animal kingdom with some representatives from single-celled or colonial eukaryotes. **C.** shows the three-pronged workflow of sequence alignment, structural alignment, and clinical interrogations. These culminate to narrow and focus on a few well-conserved amino acids that could be key to understanding the generalizable mechanism of each of the different groupings. Here, we chose three different groupings - voltage-gating, ion selectivity, and domain-swapping.

The biophysical diversity of these channels can be categorized in three major ways: preferred stimulus, ion selectivity, and domain-swapping architecture. To investigate the primary mode of stimulation, we will compare the VSD sequences and structures between voltage-gated and non-voltage-gated channels. To approach ion selectivity, we have compiled the PGD sequences and structures between potassium-selective, sodium-selective, calcium-selective, and cation-selective channels. With increases in structural knowledge of these channels within the past 20 years, certain channels have been discovered to be domain-swapped PGDs and VSDs whereas other channels are not. Here, we will speculate on some hypotheses by comparing the VSD and PGD sequences and structures of domain-swapped versus non-domain-swapped channels. Ultimately, we hope to propose testable hypotheses and, more importantly, generate a cohesive database that can be utilized to elucidate generalizable structural and functional mechanisms in this diverse channel family.

## Results

As we drew hypotheses broadly across many channels, we wanted to ensure broad representation across the animal phylogenetic tree. Hence, we compiled sequences from the National Center for Biotechnology Information (NCBI; Altschul et al., 1990; Rangwala et al., 2021) of 20 different channels (Fig. 1A) from 24 different species (Fig. 1B), ranging from eukaryotic choanoflagellates to humans (Suppl. Table 1). This ensured that the conservation we observe across channels is most likely due to function or structure rather than recent shared evolutionary ancestry. In our structural depictions, we have chosen a single structure per channel available from Protein DataBank (PDB; Berman et al., 2000), and channels without a solved structure were modeled with AlphaFold2 (Jumper et al., 2021; Mirdita et al., 2022; Steinegger and Söding, 2017; Eastman et al., 2017; Suppl Table 2). In addition to corroborating the highly conserved amino acids with existing functional studies, we cross-referenced these amino acids against the clinical variants data (ClinVar) to further draw speculations on the roles of highly conserved, yet understudied amino acids (Fig. 1C; Suppl. Tables 3-10 and Files 9-16; Landrum et al., 2014).

### Voltage-Gating

The charged residues amongst S2, S3, and S4 are well known to be responsible for voltage- sensing in voltage-sensitive channels. The negative charges in S2 and S3 stabilize the mobile positive charges of S4 as they pass through the charge-transfer center that manifests as a central aromatic on S2 (Seoh et al., 1996; Papazian et al., 1995; Aggarwal and MacKinnon, 1996; Tiwari-Woodruff et al., 1997, 2000; Starace and Bezanilla, 2004; Bell et al., 2004; Pless et al., 2011; Nilsson et al., 2022; Lacroix and Bezanilla, 2011; Barros et al., 2020; Tao et al., 2010; Ma et al., 2006; Nguyen and Horn, 2002; Gosselin-Badaroudine et al., 2012; Wu et al., 2010; Yang and Horn, 1995; Yang et al., 1996; Ma et al., 2009; Hering et al., 2018; Monteleone et al., 2017; Bezanilla, 2000). These positions are well conserved across the voltage-gated channels (Figs. 2A&B, 3A&B & 4A&B; Suppl. File 1-3). However, we observe interesting amino acid conservations that do not fit into this well-described and well-studied mechanism.

**Figure 2:**
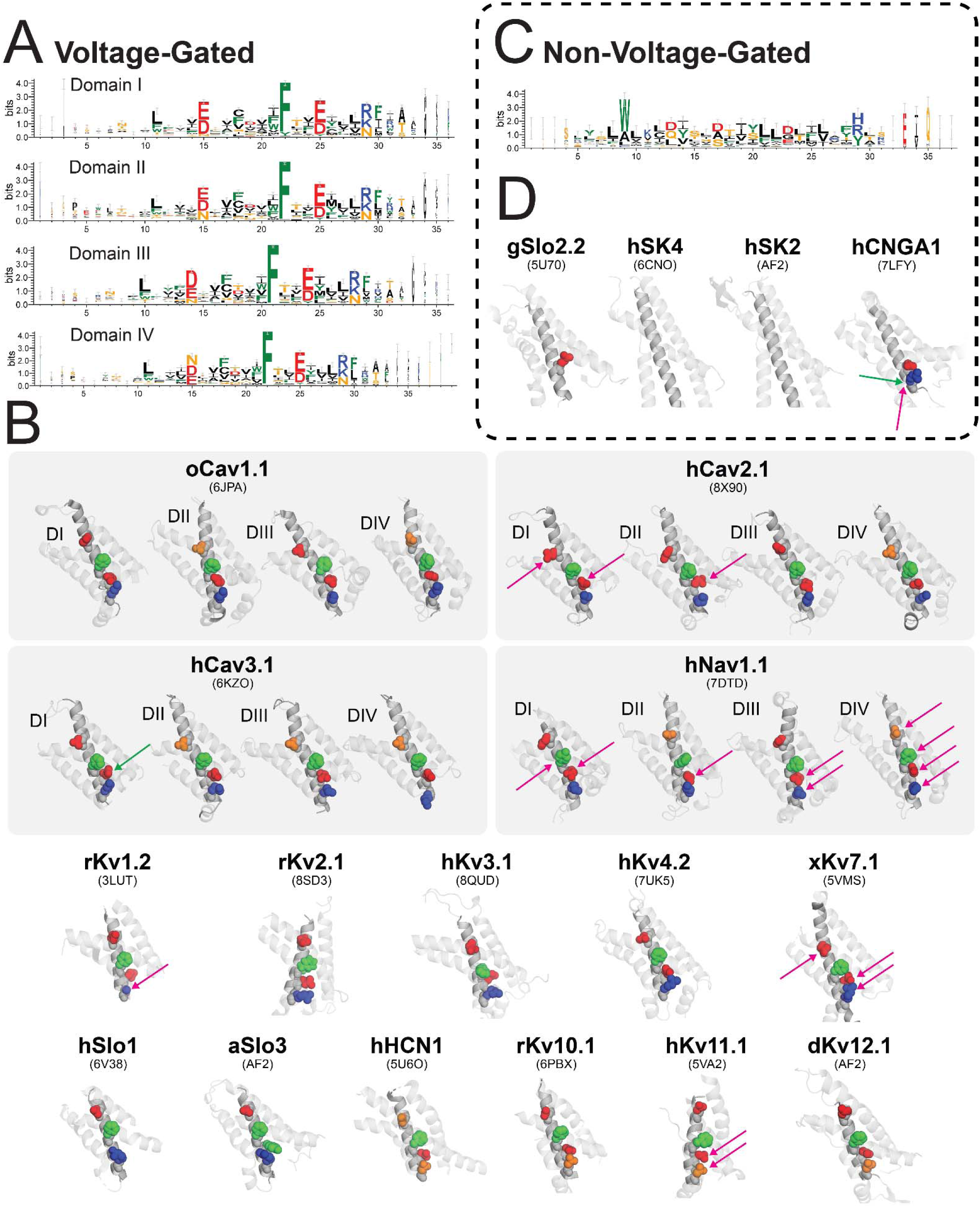
Conservation of S2 across Voltage-Gated and Non-Voltage-Gated channels. In all annotations, amino acids are color coded based on their general chemical characteristics: black - small, hydrophobic (GAPVILM); green - aromatic (FWY); orange - polar (STNQ); blue - positively charged (RKH); red - negatively charged (ED). **A.** and **C.** show the sequence logos of channels that are either voltage-gated (**A**) or non-voltage-gated (**C**). Because 24TMS channels were in the voltage-gated grouping, separate alignments were performed for each domain as a pseudo-monomer. The numbering is arbitrary and begins at the more liberal interpretation of the S2 sequence across all species variants of all channels in a particular grouping; the numbering based on each specific channel can be found in Suppl. File 1. Notice the distinct conservation of the negative residues, central phenylalanine, and positive residue in the voltage-gated channels across all domains that do not appear in the non-voltage-gated channels. **B.** and **D.** annotate the highly conserved S2 amino acids on top of the structure for each channel. Channels with solved structures are labeled with the relevant PDB code; channels without solved structures were predicted with AlphaFold2 (AF2). The arrows show the locations of pathogenic/likely pathogenic (magenta) or benign/likely benign (green) at highly conserved S2 residues. Banded arrows show the presence of both pathogenic/likely pathogenic and benign/likely benign mutations at the annotated position. The actual clinical variants can be viewed in Suppl. File 9.

#### S2 (Fig. 2; Suppl. File 1)

An S2 positively charged or polar amino acid is almost universally conserved in voltage-gated channels and located at the intracellular side of S2 (Fig. 2A&B; R - 44-45%, K - 26%, N - 27%). In solved channel structures, it is proximal to the preceding negative charge and could form a salt-bridge interaction (Fig. 2B). Furthermore, clinical mutations at this position in Kv1, 7, 11, and Nav1.1 channels are uniformly labeled as either pathogenic or likely pathogenic (Fig. 2C; 100%; Suppl. File 9). By contrast, this conservation is not well preserved in non-voltage-gated channels (Fig. 2C&D; A - 19%, R - 17%, V - 17%) and does not necessarily result in pathogenic or likely pathogenic clinical dispositions (Fig. 2D; 75%; Suppl. File 9). This position has been studied and was suggested to be a gating charge that moves across the electric field in response to voltage changes or have severe perturbations to other voltage-gating related mechanisms (Ma et al., 2006, 2009), but has remained mostly understudied - its role still unclear amidst its strong conservation and clinical relevance.

#### S3 (Fig. 3; Suppl. File 2)

The negative charge on S3 is almost uniformly an aspartate rather than a glutamate in voltage- gated channels (Fig. 3A&B; D - 99-100%) and all clinical mutations are classified as pathogenic or likely pathogenic (Fig. 3B; 100%; Suppl. File 10). This conservation is similarly not well preserved in non-voltage-gated channels. However, a preference for a negative charge is apparent (Fig. 3C&D; C - 37%, D - 35%, E - 25%) and only one mutant has been reported at this position and is likely pathogenic, though no clinical condition was cited. What is the reason for this near-uniform aspartate conservation in voltage-gated channels - could the selective pressure against glutamate be a steric effect? This position has been studied before, but only to illustrate its stabilization of the positive S4 charges (Seoh et al., 1996; Tiwari-Woodruff et al., 1997, 2000; Papazian et al., 1995; Ma et al., 2006) and not specifically the strict conservation of the aspartate over the glutamate. Given its near-uniform conservation and clinical relevance, understanding the conservation of this aspartate over glutamate is likely key to understanding the broad mechanism of voltage-sensing across voltage-gated channels.

**Figure 3:**
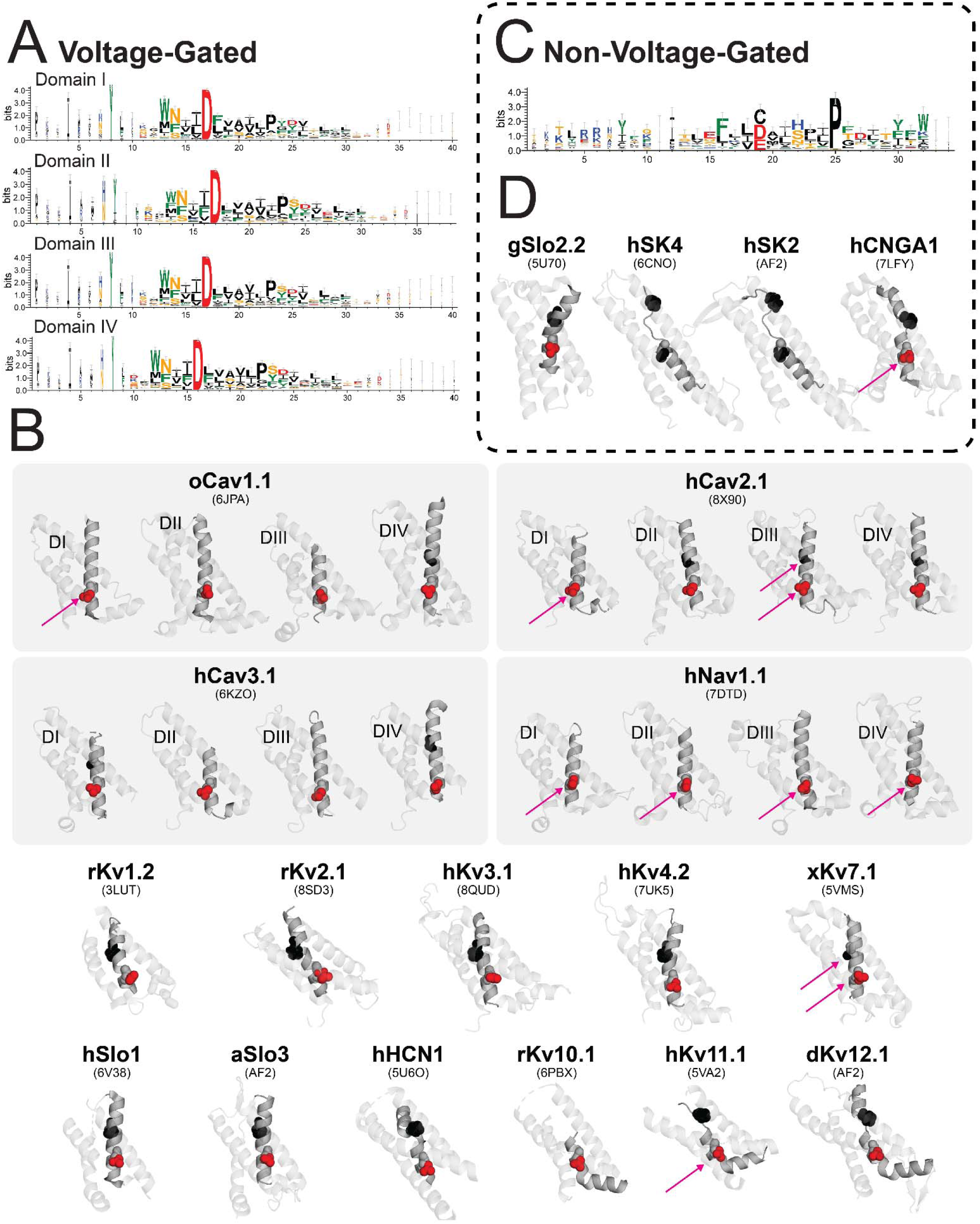
Conservation of S3 across Voltage-Gated and Non-Voltage-Gated channels. In all annotations, amino acids are color coded based on their general chemical characteristics: black - small, hydrophobic (GAPVILM); red - negatively charged (ED). **A.** and **C.** show the sequence logos of channels that are either voltage-gated (**A**) or non-voltage-gated (**C**). Because 24TMS channels were in the voltage-gated grouping, separate alignments were performed for each domain or pseudo-monomer. The numbering is arbitrary and begins at the more liberal interpretation of the S3 sequence across all species variants of all channels in a particular grouping; the numbering based on each specific channel can be found in Suppl. File 2. Notice the distinct conservation of the negative residues and the canonical S3a-b helix-breaking proline in both the voltage-gated channels and in the non-voltage-gated channels. **B.** and **D.** annotate the highly conserved S3 amino acids on top of the structure for each channel. Channels with solved structures are labeled with the relevant PDB code; channels without solved structures were predicted with AlphaFold2 (AF2). The arrows show the locations of pathogenic/likely pathogenic (magenta) at highly conserved S3 residues. The actual clinical variants can be viewed in Suppl. File 10.

Whereas the S2 positive charge and the S3 aspartate are well conserved, but understudied, there have been notable studies focusing on the S3 helix-breaking proline in 6TMS channels (Li-Smerin and Swartz, 2001) (Fig. 3A&B; P - 61-64%). These studies highlight that this proline is key to the mobility of the latter portion of S3 (S3b) in the voltage-gating mechanism. However, this proline is not present in many voltage-gated calcium and sodium channels and is neither well conserved across the 24TMS channels nor across domains (I-IV). Surprisingly, non-voltage-gated channels conserve this proline at an even higher rate (Fig. 3C&D; P - 94%). This draws the question of how broadly necessary a mobile S3b is in the overall mechanism of voltage-gating and whether the mobility of S3b is necessary only in the 6TMS channels.

#### S4 (Fig. 4; Suppl. File 3)

S4 is home to many positively charged amino acids that move across the membrane electric field in response to membrane voltage changes. These amino acids are known as gating charges and are critical to voltage-sensing in voltage-gated channels. Generally, the first three positively charged residues are cited to be the most critical gating charges (Bell et al., 2004; Papazian et al., 1995; Aggarwal and MacKinnon, 1996; Hering et al., 2018; Ma et al., 2006; Seoh et al., 1996; Yang and Horn, 1995; Yang et al., 1996). We see high conservation of all the positively charged amino acids in our alignments as well (Fig. 4A&B; R - 33-97%, K - 7-35%). In particular, the central canonically named R4 position is nearly uniformly conserved to be an arginine greater than 95% (Fig. 4A&B). The immutability of this arginine instead of a lysine is interesting. In Aggarwal & MacKinnon, 1996, Kv1 channels with this R to K substitution were noted to be voltage-sensitive but no Q-V curves were shown (Aggarwal and MacKinnon, 1996). Clinical mutants at this position are cited to have pathogenic or likely pathogenic dispositions (Fig. 4B; 97.6%; Suppl. File 11). There is a likely pathogenic clinical mutant resulting in an arginine to lysine substitution in domain IV of Cav1.1 (Fig. 4B; Suppl. File 11), but no work has been done to biophysically characterize this substitution. Clearly, the positive charge at this position is critical to voltage-gating, but the biophysical consequence or evolutionary pressure to conserve an arginine over a lysine is unclear.

**Figure 4:**
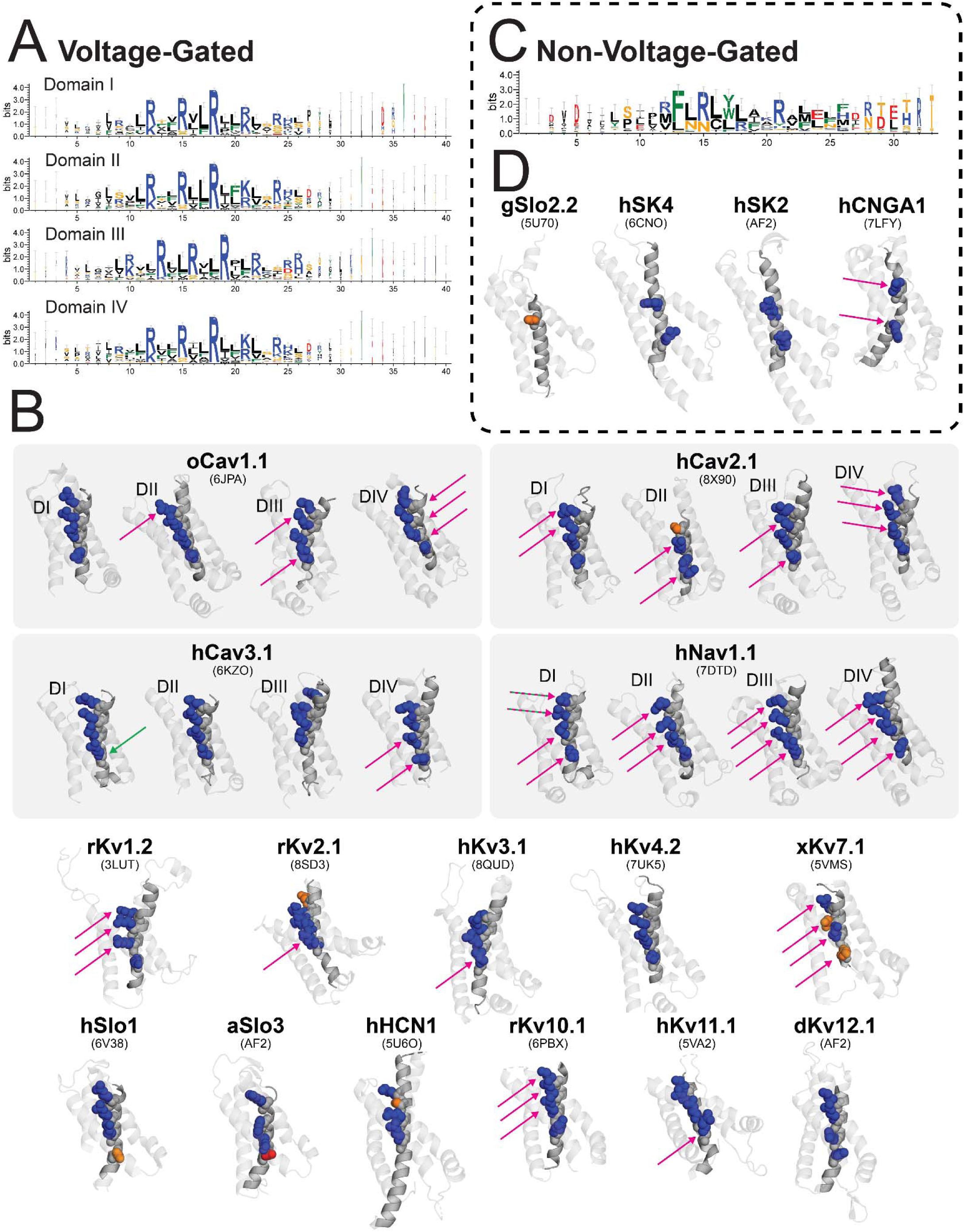
Conservation of S4 across Voltage-Gated and Non-Voltage-Gated channels. In all annotations, amino acids are color coded based on their general chemical characteristics: orange - polar (STNQ); blue - positively charged (RKH); red - negatively charged (ED). **A.** and **C.** show the sequence logos of channels that are either voltage-gated (**A**) or non-voltage-gated (**C**). Because 24TMS channels were in the voltage-gated grouping, separate alignments were performed for each domain or pseudo-monomer. The numbering is arbitrary and begins at the more liberal interpretation of the S4 sequence across all species variants of all channels in a particular grouping; the numbering based on each specific channel can be found in Suppl. File 3. Notice the distinct conservation of the four positive residues in the voltage-gated channels but only two positive residues in the non-voltage-gated channels. **B.** and **D.** annotate the highly conserved S4 amino acids on top of the structure for each channel. Channels with solved structures are labeled with the relevant PDB code; channels without solved structures were predicted with AlphaFold2 (AF2). The arrows show the locations of pathogenic/likely pathogenic (magenta) and benign/likely benign (green) at highly conserved S4 residues. Banded arrows show the presence of both pathogenic/likely pathogenic and benign/likely benign mutations at the annotated position. The actual clinical variants can be viewed in Suppl. File 11.

Interestingly, in non-voltage-gated channels, we also observe high conservation of two arginines (Fig. 4C&D both R - 69%) relating the canonically denoted R2 and R4 positions. Clinical SNPs at these two positions all occur in CNGA3 channels and are cited to be pathogenic or likely pathogenic (Fig. 4D; 100%; Suppl. File 11). The conservation at these positions in non-voltage-gated channels is unknown. Whether this conservation is due to functional importance, structural stability, or as an evolutionary vestige is unclear.

### Ion Permeation

The ion selectivity of the VGL superfamily channels is primarily affected by the amino acids of the reentrant pore helix that canonically form the selectivity filter. Here, we will highlight some of the key amino acids of the selectivity filter that have been found to be responsible for ion selectivity (Wu et al., 2022; Senatore et al., 2013; Yang et al., 1997; Coonen et al., 2022; Bernsteiner et al., 2017; Kopec et al., 2019; Abderemane-Ali et al., 2019; Ellinor et al., 1995; Yang et al., 1993; Parent and Gopalakrishnan, 1995; Tsushima et al., 1997; Favre et al., 1996; Chiamvimonvat et al., 1996; Heinemann et al., 1992; Morais-Cabral et al., 2001; Doyle et al., 1998).

#### Pore (Fig. 5; Suppl. File 4)

The latter portion of the pore helix is typically structurally resolved to be pore-facing, resulting in interactions with the ions that permeate through the pore, canonically called the selectivity filter. In potassium channels, these positions are well-conserved as a GYG or a GFG motif (Doyle et al., 1998; Morais-Cabral et al., 2001; Bernsteiner et al., 2017; Kopec et al., 2019); Fig. 5A&B; G - 99.6%, Y/F - 70.4%/28.8%, G - 99.6%) and mutations in these positions are unequivocally pathogenic or likely pathogenic (Fig. 5B; 100%). Calcium and sodium channels conserve a different set of motifs, befitting their calcium and sodium selectivities. Sodium channels have a particular set of inward-facing amino acids that interact with the permeating sodium and form an asymmetric selectivity filter (Fig. 5C&D; DI - D - 100%, DII - E - 95%, DIII - K - 74%, DIV - A - 95%). All mutations that occur at these positions are pathogenic or likely pathogenic (Fig. 5D). In calcium channels, each domain’s pore helix donates a pore-facing negative charge (Fig. 5E&F; E - 65-97%, D - 35-0%) that interacts with the permeating calcium ions (Heinemann et al., 1992; Parent and Gopalakrishnan, 1995; Yang et al., 1993; Ellinor et al., 1995; Abderemane-Ali et al., 2019). All mutations at these negative charges result in pathogenic or likely pathogenic clinical dispositions (Fig. 5F; Suppl. File 12).

**Figure 5:**
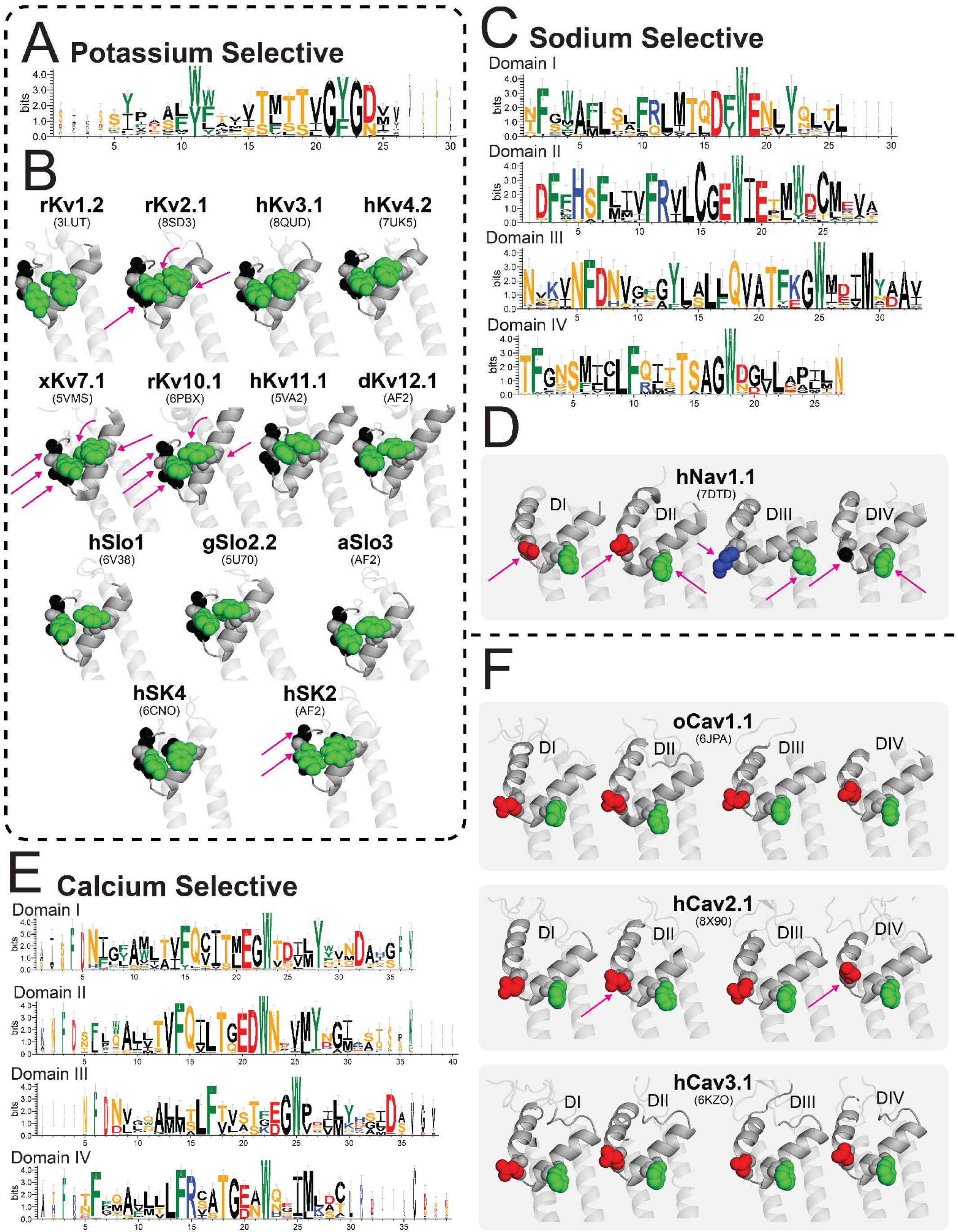
Conservation of Pore Helix across different ion selectivities. In all annotations, amino acids are color coded based on their general chemical characteristics: black - small, hydrophobic (GAPVILM); green - aromatic (FWY); orange - polar (STNQ); blue - positively charged (RKH); red - negatively charged (ED). **A.**, **C.**, and **E.** show the sequence logos of potassium (**A**), sodium (**C**), and calcium (**E**). Because 24TMS channels were in sodium and calcium selective groupings, separate alignments were performed for each domain or pseudo-monomer. The numbering is arbitrary and begins at the more liberal interpretation of the pore helix sequence across all species variants of all channels in a particular grouping; the numbering based on each specific channel can be found in Suppl. File 4. Here, the selectivity filter amino acids (GYG for potassium, DEKA for sodium, and E/D for calcium) are well conserved within each grouping. Additionally, aromatics preceding the selectivity filter are also very well conserved. **B.**, **D.** and **F.** annotate the highly conserved pore helix amino acids on top of the structure for each channel. Channels with solved structures are labeled with the relevant PDB code; channels without solved structures were predicted with AlphaFold2 (AF2). The arrows show the locations of pathogenic/likely pathogenic (magenta) at highly conserved pore helix residues. Curved arrows annotate the positions that are not visible in this view and are the amino acids that face into the page. The actual clinical variants can be viewed in Suppl. File 12.

The beginning portion of the pore helix is critically important to permeation as well (Yang et al., 1997; Wu et al., 2022; Coonen et al., 2022). Potassium channels have well-conserved aromatics (Fig. 5A&B) that are also present in calcium and sodium channels. Interestingly, a tryptophan to phenylalanine mutation can rend Kv1 channels impermeant to any ions - functionally closing the pore, structurally dilating the selectivity filter (Yang et al., 1997; Wu et al., 2022; Coonen et al., 2022).. Well-conserved aromatics are also seen in the different domains of the 24TMS channels (Fig. 5C-F); it is unclear whether these positions also hold similarly critical roles in allowing for permeation through the selectivity filter.

### Domain-Swapping

The functional consequence of the domain-swapping phenomenon between VSDs and PGDs of specific ion channels is still unknown. Only hints are provided of its functional biophysical consequence. For example, speculatively, the VSD-PGD domain-swapping could be the mechanism by which a two-step voltage-gating process might occur. KCNQ channels are driven by a two-step voltage-gating process - the VSD interacts with its sequence-contiguous PGD for one step, and with its domain-swapped neighboring PGD for the other step (Hou et al., 2020). Here, we aim to highlight some well-conserved amino acids across domain-swapped and non-domain-swapped ion channels. Undoubtedly well-conserved amino acids could serve multiple roles. However, to highlight understudied positions that could be critical for domain-swapping in particular, we will ignore the well-conserved amino acids that have been cited in the previous sections.

#### S2 (Fig. 6; Suppl. File 5)

Aside from the negative and central aromatic conserved amino acids that were also observed in the alignments of the voltage-gating divisions, we also observe a high conservation of a small polar amino acid (Fig. 6A&B; T - 46.4-57%, S - 9.3-17%) in domain-swapped channels. This conservation is not observed in non-domain-swapped channels (Fig. 6C&D; L - 47%, I - 24.4%). There are only two pathogenic or likely pathogenic mutations in Nav1.1 that have been reported but have not been biophysically studied (Fig. 6B; Suppl. File 9; Margherita Mancardi et al., 2006). However, given that S2 is distant from the VSD-PGD interaction sites, it is unclear whether this position is truly critical for domain-swapping or if it has another functional or structural role.

**Figure 6:**
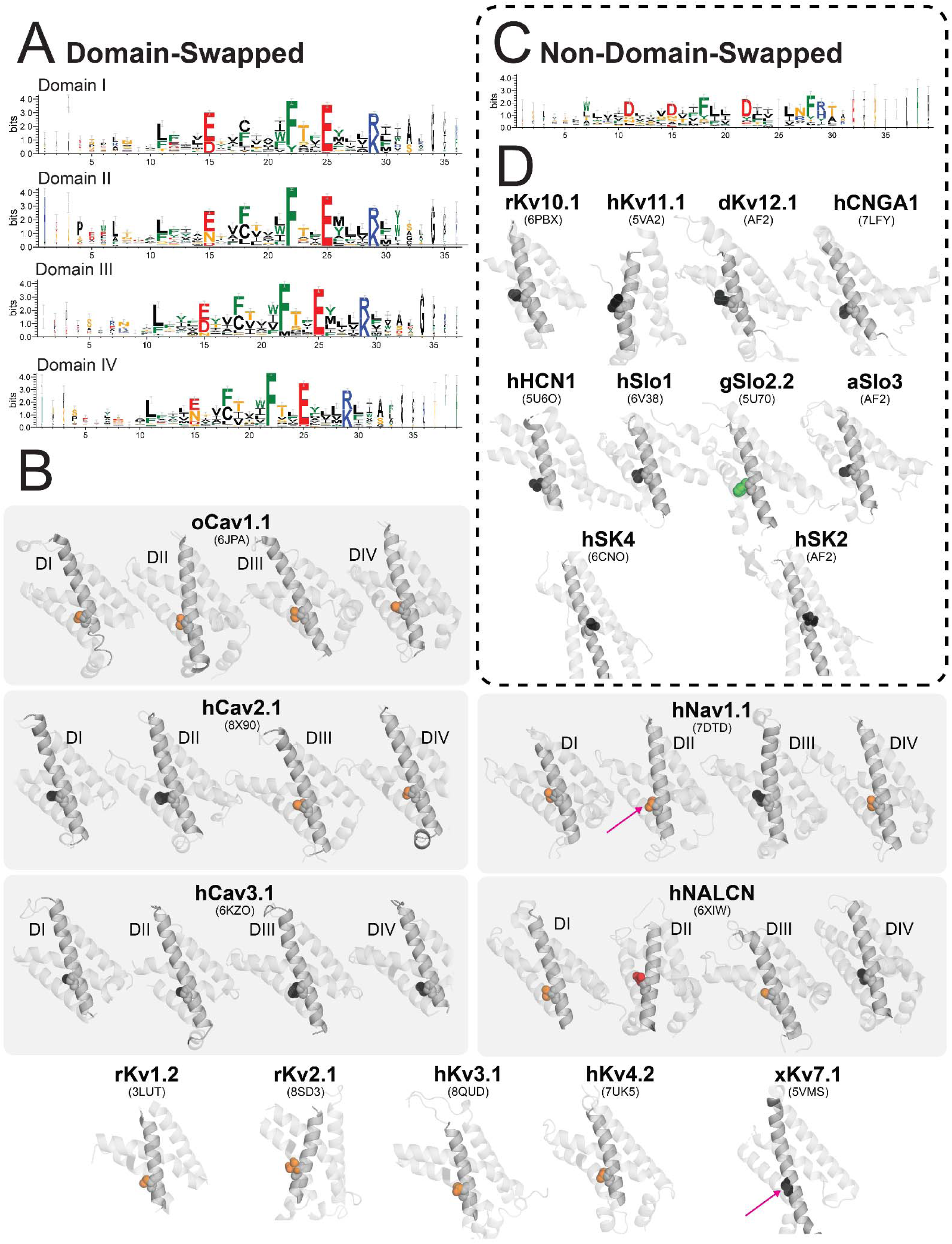
Conservation of S2 across different domain-swapping architectures. In all annotations, amino acids are color coded based on their general chemical characteristics: black - small, hydrophobic (GAPVILM); green - aromatic (FWY); orange - polar (STNQ); blue - positively charged (RKH); red - negatively charged (ED). **A.** and **C.** show the sequence logos of domain-swapped (**A**) and non-domain-swapped (**C**) channels. Because all 24TMS channels were in domain-swapped groupings, separate alignments were performed for each domain or pseudo-monomer. The numbering is arbitrary and begins at the more liberal interpretation of the pore helix sequence across all species variants of all channels in a particular grouping; the numbering based on each specific channel can be found in Suppl. File 5. Here, a single polar amino acid is conserved within the domain-swapped channels but is highly conserved to be a hydrophobic amino acid in non-domain-swapped channels. This amino acid sits directly following the phenylalanine charge transfer center in voltage-gated channels (Fig. 2). **B.** and **D.** annotate the highly conserved amino acids on top of the structure for each channel that have not been shown to be responsible for voltage-gating. Channels with solved structures are labeled with the relevant PDB code; channels without solved structures were predicted with AlphaFold2 (AF2). The arrows show the locations of pathogenic/likely pathogenic (magenta) at highly conserved pore helix residues. The actual clinical variants can be viewed in Suppl. File 9.

#### S3 (Fig. 7; Suppl. File 6)

Similarly to the negative and aromatic amino acids of S2, the negatively charged aspartate is highly conserved across both the non-domain-swapped and domain-swapped channels. Aside from this aspartate, there are two positions of note.

**Figure 7:**
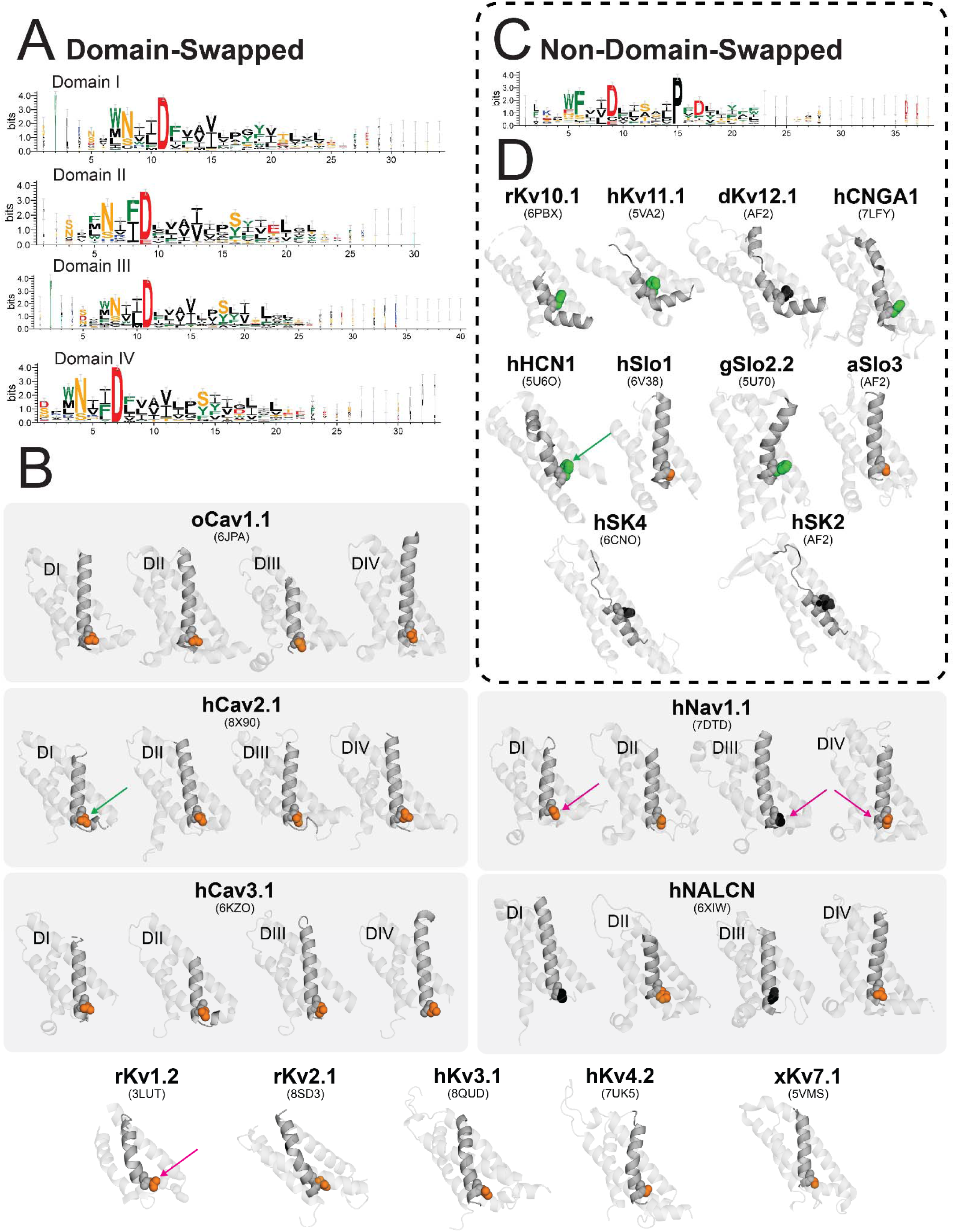
Conservation of S3 amino acids across different domain-swapping architectures. In all annotations, amino acids are color coded based on their general chemical characteristics: black - small, hydrophobic (GAPVILM); green - aromatic (FWY); orange - polar (STNQ); blue - positively charged (RKH); red - negatively charged (ED). **A.** and **C.** show the sequence logos of domain-swapped (**A**) and non-domain-swapped (**C**) channels. Because all 24TMS channels were in domain-swapped groupings, separate alignments were performed for each domain or pseudo-monomer. The numbering is arbitrary and begins at the more liberal interpretation of the pore helix sequence across all species variants of all channels in a particular grouping; the numbering based on each specific channel can be found in Suppl. File 6. Here, a single polar amino acid is conserved within the domain-swapped channels but is not well conserved in non-domain-swapped channels. **B.** and **D.** annotate the highly conserved amino acids on top of the structure for each channel that have not been shown to be responsible for voltage-gating. Channels with solved structures are labeled with the relevant PDB code; channels without solved structures were predicted with AlphaFold2 (AF2). The arrows show the locations of pathogenic/likely pathogenic (magenta) and benign/likely benign at highly conserved pore helix residues. The actual clinical variants can be viewed in Suppl. File 10.

In domain-swapped channels, a polar or histidine amino acid is highly conserved (Fig. 7A&B; (Fig. 7A&B; N - 60.4-78.4%, S - 15.8-20.2%, H - 0-8%). This strong conservation in domains II and IV of domain-swapped channels lies in stark contrast to the conservation seen in non-domain-swapped channels, wherein a phenylalanine is conserved (Fig. 7C&D; F - 68.9%, S - 13%). A single mutation at this position occurs in patients and is likely pathogenic (Fig. 7B; Suppl. File 10). Given its high conservation between the two groups, it is likely that this position holds significance in these channels. However, the role of this position in determining domain-swapping or in some other functional capacity remains understudied.

#### S5 (Fig. 8; Suppl. File 7)

Amino acids of S5 have been relatively understudied compared to the well-conserved amino acids of other transmembrane segments. However, given that they are connected to the preceding S4 by a linker region and likely have interactions with S4, there are likely well-conserved amino acids that might play a role in the different VSD-PGD interactions between domain-swapped and non-domain-swapped channels. We hereby observe a couple of highly conserved residues in both the domain-swapped and non-domain-swapped that might hold some functional relevance.

**Figure 8:**
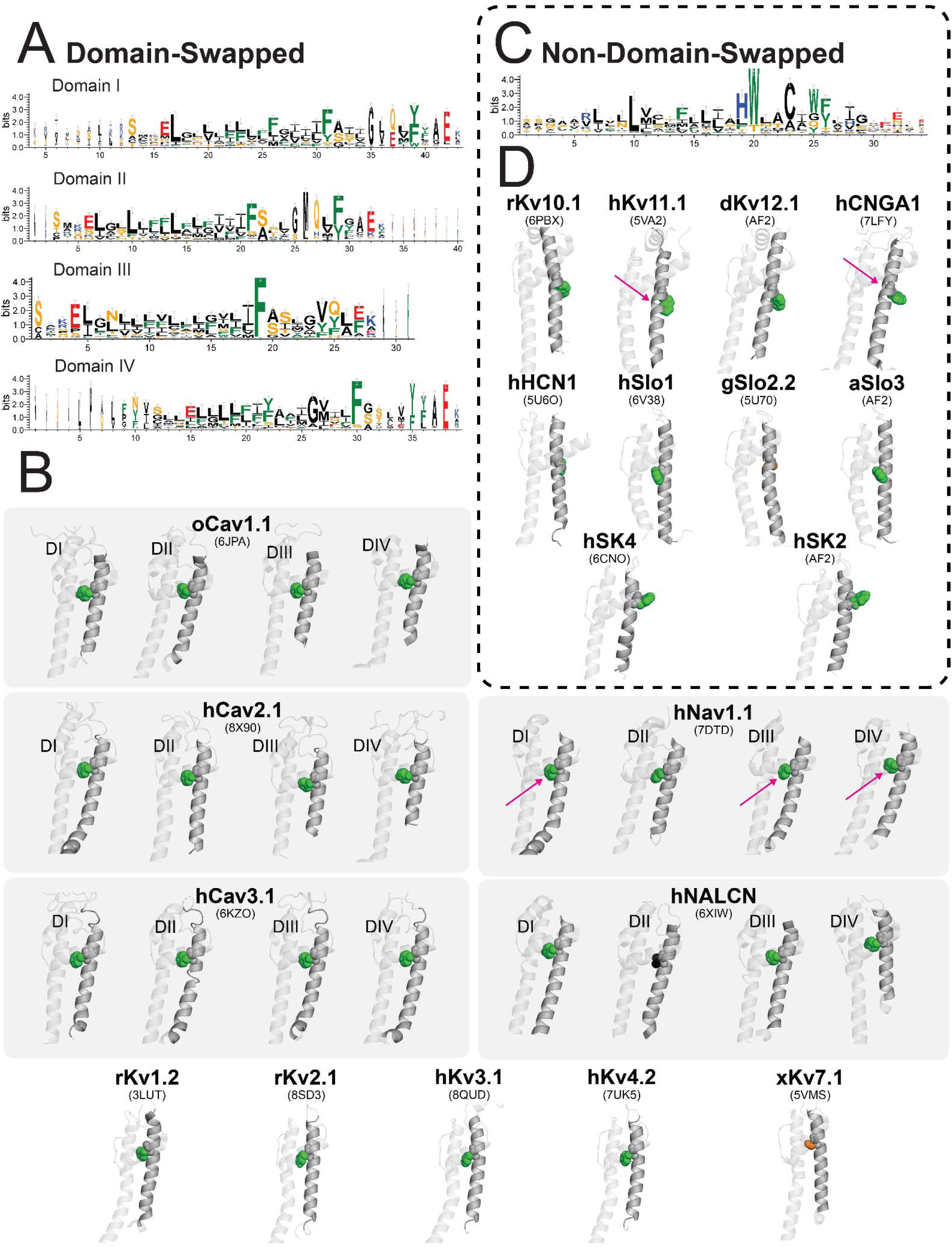
Conservation of S5 amino acids across different domain-swapping architectures. In all annotations, amino acids are color coded based on their general chemical characteristics: black - small, hydrophobic (GAPVILM); green - aromatic (FWY); orange - polar (STNQ); blue - positively charged (RKH); red - negatively charged (ED). **A.** and **C.** show the sequence logos of domain-swapped (**A**) and non-domain-swapped (**C**) channels. Because all 24TMS channels were in domain-swapped groupings, separate alignments were performed for each domain or pseudo-monomer. The numbering is arbitrary and begins at the more liberal interpretation of the pore helix sequence across all species variants of all channels in a particular grouping; the numbering based on each specific channel can be found in Suppl. File 7. **B.** and **D.** annotate the highly conserved amino acids on top of the structure for each channel that have not been shown to be responsible for voltage-gating. Channels with solved structures are labeled with the relevant PDB code; channels without solved structures were predicted with AlphaFold2 (AF2). The arrows show the locations of pathogenic/likely pathogenic (magenta) and benign/likely benign at highly conserved pore helix residues. The actual clinical variants can be viewed in Suppl. File 13.

In non-domain-swapped channels, there is a central tryptophan that is highly conserved (Fig. 8C&D; W - 84.5%, T - 10.7%). This position is similar in location to the well-conserved non-tryptophan aromatic in domain-swapped channels (Fig. 8A&B; F - 36.5-97.0%, Y - 0-37.9%). Strangely in domain-swapped channels, this position is facing inwards towards the pore rather than outwards towards the VSD (Fig. 8B); whereas in non-domain-swapped channels, this position is sometimes facing inwards (Slo family) and outwards other times (CNBD family and SK family; Fig. 8D). Interestingly, the two clinical mutations in non-domain-swapped channels at this position are posited to be likely pathogenic (Fig. 8D). Similarly, there are two clinical mutations in domain-swapped channels that are also posited to be likely pathogenic as well (Fig. 8B; Suppl. File 13). It is unclear whether this structural and sequence mismatch in alignment is due to a misalignment or carries structure-function implications.

#### S6 (Fig. 9; Suppl. File 8)

S6 is a pore-lining helix where the gate that opens and closes in response to the appropriate stimulus resides. However, the location of the gate does not seem to be generalizable; in some cases, there is a bundle-crossing, whereas other gates are within the deep-pore region (Zhou et al., 2011; Chen et al., 2014; Chen and Aldrich, 2011; Imbrici et al., 2009; Whicher and Mackinnon, 2019; Xue et al., 2021; He et al., 2022; Liu et al., 2023). We will highlight some amino acids that have been well-described functionally and illuminate other conserved amino acids whose roles are not well elucidated in all channels.

**Figure 9:**
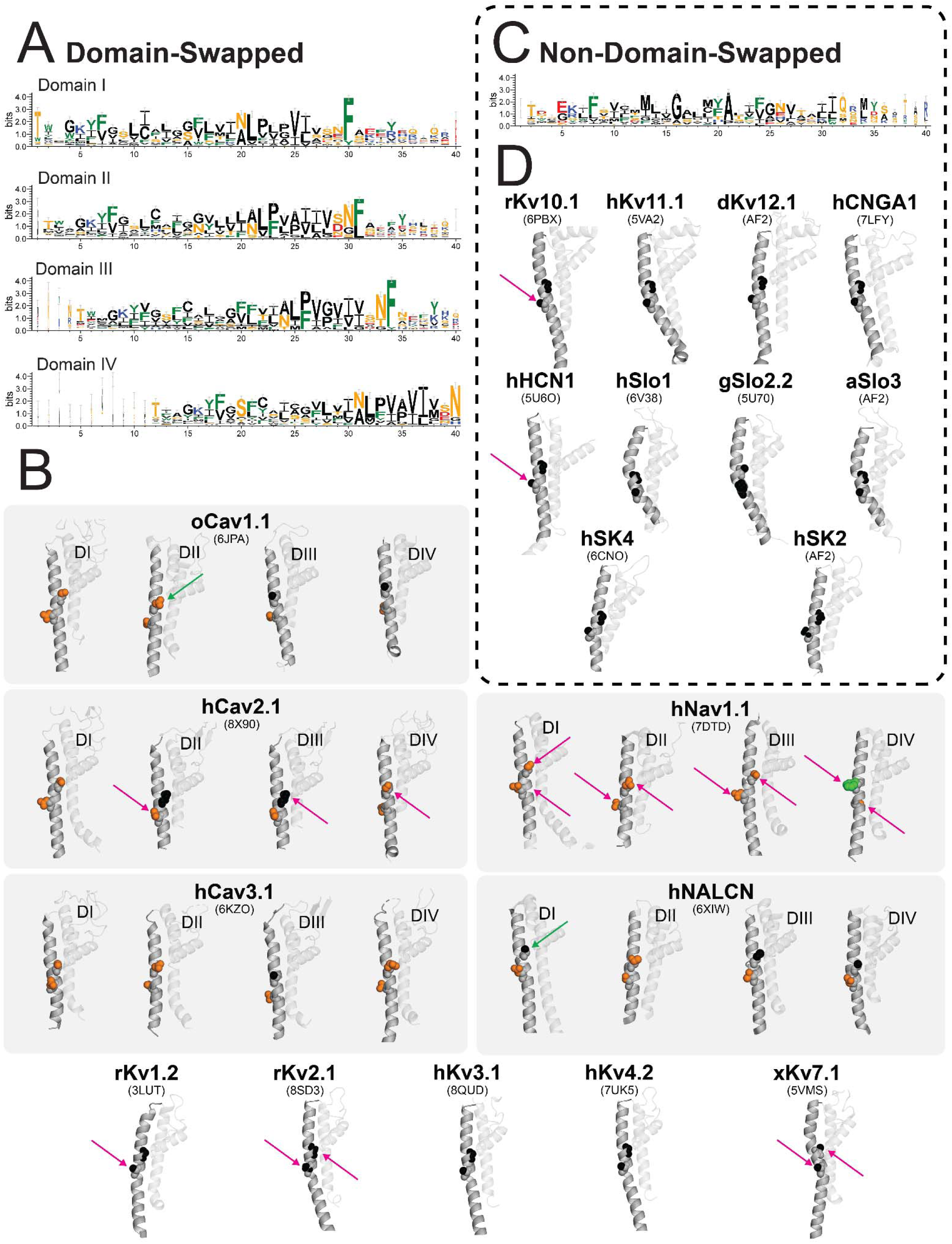
Conservation of S6 amino acids across different domain-swapping architectures. In all annotations, amino acids are color coded based on their general chemical characteristics: black - small, hydrophobic (GAPVILM); green - aromatic (FWY); orange - polar (STNQ); blue - positively charged (RKH); red - negatively charged (ED). **A.** and **C.** show the sequence logos of domain-swapped (**A**) and non-domain-swapped (**C**) channels. Because all 24TMS channels were in domain-swapped groupings, separate alignments were performed for each domain or pseudo-monomer. The numbering is arbitrary and begins at the more liberal interpretation of the pore helix sequence across all species variants of all channels in a particular grouping; the numbering based on each specific channel can be found in Suppl. File 8. In particular, note the high conservation of glycine and alanine in the central area of the non-domain-swapped channels. Commonly this position is conserved to a polar or an alanine in domain-swapped channels. **B.** and **D.** annotate the highly conserved amino acids on top of the structure for each channel that have not been shown to be responsible for voltage-gating. Channels with solved structures are labeled with the relevant PDB code; channels without solved structures were predicted with AlphaFold2 (AF2). The arrows show the locations of pathogenic/likely pathogenic (magenta) and benign/likely benign at highly conserved pore helix residues. The actual clinical variants can be viewed in Suppl. File 14.

A glycine central to S6 is known amongst potassium channels to be a critical amino acid that allows for S6 mobility in response to gate movement (Magidovich and Yifrach, 2004; Yifrach and MacKinnon, 2002; Zhou et al., 2011; Ding et al., 2005; Chen and Aldrich, 2011; Chen et al., 2014; Hardman et al., 2007; Boulet et al., 2007); canonically, it has been labeled as the glycine hinge. We observe high conservation of this position in non-domain-swapped channels (Fig. 9C&D; G - 81.4%, A - 11.6%). Such high conservation for a glycine does not appear in domain-swapped channels (Fig. 9A&B; G - 37.3-47.3%; A - 18.5-22.4%; S - 10.9-14.5%). Whereas there are no clinical mutations at this position in non-domain-swapped channels, there are 20 clinical mutations in domain-swapped channels, of which 18 are pathogenic or likely pathogenic (Fig. 9B; 90%; Suppl. Table 10 and 16). Regardless, it is unclear and highly speculative as to whether this well-conserved glycine has a functional role in determining a non-domain-swapped configuration and whether this position has a generalizable role in domain-swapped channels, or if the high conservation is an artifact of potassium channels dominating the non-domain-swapped channels (Suppl. File 8).

A lesser-studied amino acid is the highly-conserved alanine in non-domain-swapped channels about five positions after the central S6 glycine hinge (Fig. 9C&D; A - 84.3%). This position is split as an alanine or an asparagine (Fig. 9A&B; N - 39.8-48.8%, A - 47.3-47.5%) whereby potassium channels generally conserve an alanine and other channels generally conserve an asparagine. Aside from some studies in Slo1 channels that investigate the role of this position in forming the gate (Zhou et al., 2011; Chen et al., 2014), details about the role of this alanine in other channels are sparse. Supportingly, there are 5 mutations at this position in non-domain-swapped channels that are likely to give pathogenic clinical dispositions (Fig. 9D; 100%; Suppl. File 14). Substitutions at this position can result in either constitutive open or closed channels (Zhou et al., 2011; Chen et al., 2014). Interestingly, there are 28 pathogenic or likely pathogenic mutations at this position in domain- swapped channels (Fig. 9B; 100%; Suppl. File 14). Whether this position holds a similarly critical role in other channels as it does in Slo1 channels is unclear and would warrant further investigation.

## Discussion

In our comparisons between groupings, we have observed highly conserved positions that are well-corroborated with functional or structural studies. We also revealed highly conserved residues that are less studied, have clinical indications, and are likely key to furthering our understanding of the foundational properties of each grouping. We hope that this compiled database can be used as a further hypothesis generator for the field. However, there are some key limitations to this database that must be addressed.

### Assumptions of Our Approach

Although our approach is clearly corroborated by previous functional and structural work, there are key assumptions and limitations. *Assumption 1:* We assume that conservation indicates functional or structural importance and is not derived from shared evolutionary lineage. We attempted to diffuse this critical confound of sequence alignment analyses by *A.* utilizing a broad sampling of species across the Animal Kingdom and *B.* aligning across distantly-related channels. *Assumption 2:* We assume that the same channels from distantly-related species generally carry the same fundamental functional and structural properties. It is not outside the possibility that species-specific mutations might change these fundamental properties, but we believe that having 24 species across the Animal Kingdom could help reduce the probability that conservation is occurring by chance. However, only studies that investigate the species variants in less well-studied species can test this assumption. *Assumption 3:* All shown structures are in similar “states” even across different ion channels and different methodologies. In recent years, there has been an emerging understanding that the lipid environment has a large impact on the observed states of ion channels. Generally, we chose structures that were determined to be in the open state and with the voltage-sensor in the “up” or depolarized state. This could not be done with AlphaFold2 predicted structures so we assumed them to be in the open and depolarized states. *Assumption 4:* Evolutionary change at each position is independent of all other mutations at other positions. Here, we assume no co-evolution for the sake of simplicity of analysis. However, in cases such as the charges and countercharges of S4 and S2-3, there could be coevolution that occurs. *Assumption 5*: The fundamental properties of voltage-gating, ion selectivity, and domain-swapping architecture are not determined by the intersegment linkers. Although the linker regions can influence voltage-gating and ion selectivity, the general categorization of these fundamental properties is dictated by the transmembrane segments. However, domain-swapping architecture could rely on the length of the S4-S5 linker that tethers the VSD to the PGD. Here, our approach is completely ignorant of these variabilities that could drastically affect the domain-swapping architecture.

### A Cohesive Database

Here, we propose many hypotheses and understudied positions that could have generalizable roles across the different comparisons that we have performed. By overlooking the complexities of each channel, we attempt to garner attention to the amino acids that seem foundational to generalizable fundamental properties. However, our database does not only need to be utilized to investigate the broadstroke mechanisms but also can generate hypotheses to tease apart the nuances of individual channel function and structure.

Critically, we corroborate our findings in brief with previous functional literature on voltage-gating, highlight some positions in S2 and S3 that could have an impactful generalizable role in voltage-gating, and draw questions as to how non-voltage-gated channels might have lost their voltage-sensing capabilities through various mutations at a few critical amino acids. We also find evidence that parrots the seminal works done in the selectivity filter and show that our bioinformatics methodology does indeed highlight known functional amino acid positions. Finally, we speculate amino acid positions that might play a role in the emerging domain-swapping phenomenon that occurs in the VGL superfamily. Our approach capitalizes on the breadth of bioinformatics, functional, and structural work that has been done on the channels of the VGL superfamily to generate a database from which hypotheses of broad or detailed scopes can be drawn and tested.

## Supporting information

All referenced PDB files

## Acknowledgements

We would like to thank past members of the Aldrich Lab for their continued support in this endeavor. In particular, we acknowledge the astute contributions of Thomas Middendorf, PhD and D. Brent Halling, PhD. Additionally, we thank the fruitful discussions with the ion channel group at University of Texas at Austin with special thanks to Harold Zakon, PhD, Eric Senning, PhD, Marcel Goldschen-Ohm, PhD, and Jonathan Pierce, PhD. We thank Julia York, PhD as well for insightful discussions and pointing us in the right directions. We also would like to thank the Jegla Lab and Benjamin Simonson, PhD for providing certain *M. leidyi* sequences and for continued highlighting of the pitfalls of our endeavor. We would also like to acknowledge the funding sources for this work: F31NS124283 (FY), F32DC022491 (FY), and R01GM127332 (RWA).

## Materials and Methods

### Sequence Selection Across Many Phyla (Fig. 1; Suppl. Table 1)

Channels of the VGL superfamily that formed from six-transmembrane (6TMS) homotetramers or 24 transmembrane (24TMS) monomers were chosen for this alignment. TRP channels were excluded because a similar methodology and analysis has already been done (Fig. 1A) (Huffer et al., 2020; Cabezas-Bratesco et al., 2022). The channels that were ultimately included in this alignment were K_v_1-4, K_v_ 7, K_v_ 10-12, CNG, HCN, K_Ca_1-5, Nav 1, Cav 1-3, and NALCN. We decided to ignore obligate heterotetrameric channels (Kv 5, 6, 8, 9) from this alignment. 24 species’ genomes were BLAST’ed for these channels (Altschul et al., 1990). These species are: *Capsaspora owczarzaki* (single-celled eukaryote), *Salpingoeca rosetta* (marine colonial eukaryote), *Mnemiopsis leidyi* (warty comb jelly; Simonson et al., 2024)*, Bolinopsis microptera* (comb jelly), *Amphimedon queenslandica* (sponge), *Nematostella vectensis* (starlet sea anemone), *Hydra vulgaris* (swiftwater hydra), *Helobdella robusta* (leech), *Lottia gigantea* (owl limpet), *Aplysia californica* (California sea hare), *Schistosoma mansoni* (blood fluke), *Caenorhabditis elegans* (nematode), *Drosophila melanogaster* (fruit fly), *Priapulus caudatus* (cactus worm), *Strongylocentrotus purpuratus* (purple sea urchin), *Ciona intestinalis* (vase tunicate), *Branchiostoma floridae* (Florida lancelet), *Petromyzon marinus* (sea lamprey), *Callorhinchus milii* (elephant shark), *Pogona vitticeps* (bearded dragon), *Alligator mississippiensis* (American alligator), *Gallus gallus* (chicken), *Mus musculus* (mouse), and *Homo sapiens* (human) (Fig. 1B). These species with well-annotated genomes were chosen to sample across multiple phyla. The human channel genes were used to BLAST each of the other species and the highest matching transcript with the most exons was used (Rangwala et al., 2021). The species variant was then reverse BLAST’ed into the *Drosophila* genome to ensure the correct identification of the species variant of the channel; following, a reverse BLAST into the whole NCBI database was performed to verify that these species variants were most likely from the species of interest (Fig. 1C). The FASTA codes used can be seen in Suppl. Table 1.

### Selection and Alignment of Structures (Fig. 1C; Suppl. Table 2)

Structures were compiled from PDB (Supp Table 2) and visualized in PyMol (Schrödinger, LLC, 2015; Berman et al., 2000). An open, depolarized channel was utilized for the structural alignment, with the exception of HCN whereby two structures - an open channel (PDB: 8T50) and a depolarized channel (PDB: 5U6O) - structures were used separately. Each structure was used to confirm the amino acids of each transmembrane segment to allow for alignments across each transmembrane segment across all channels (Fig. 1C; Tao and MacKinnon, 2019; Lee and MacKinnon, 2018; Hite and MacKinnon, 2017; Chen et al., 2010; Fernández-Mariño et al., 2023; Liang et al., 2024; Ye et al., 2022; Sun and MacKinnon, 2017; Whicher and Mackinnon, 2019; Wang and MacKinnon, 2017; Lee and MacKinnon, 2017; Burtscher et al., 2024; Pan et al., 2021; Zhao et al., 2019b; Li et al., 2024; Zhao et al., 2019a; Kschonsak et al., 2020).

Each channel structure was parsed into two parts: the voltage-sensitive domain (VSD) consisting of TMS 1-4, and the pore domain (PGD) consisting of TMS 5-6. This was done to prevent misalignments due to domain-swapping derived variability in PGD-VSD positioning. Both domains were aligned as monomers. K_Ca_1 was used as the node for structural alignments of all 6TMS channels with two exceptions. The structural alignment of SK has K_v_ 7 as its node. The PGD of the hyperpolarized, open state of HCN (Burtscher et al., 2024) was used to align with the PGD of other channel structures. The VSD-depolarized, closed state of HCN (Lee and MacKinnon, 2017) was used to align the VSD with other depolarized VSD structures. Almost all of the PGDs and VSDs of Nav 1 were structurally aligned to KCa1. Nav 1 DII Pore was the only exception to this with K_v_1 Pore as its node. The VSDs and PGDs of Cav 1-3 and NALCN were then aligned to their DI Nav 1 equivalent.

The highly conserved amino acid positions that were noted in the Alignment of Transmembrane Segments were highlighted and investigated in the structure. The color coding is as follows: black - small, hydrophobic (GAPVILM); green - aromatic (FWY); orange - polar (STNQ); blue - positively charged (RKH); red - negatively charged (ED).

Channels with no available structure in the RCSB Protein Data Bank were predicted with AlphaFold2 (Jumper et al., 2021) using the ColabFold-AF2 platform (Mirdita et al., 2022). Sequence tetramers of *Danio rerio* K_v_12.2, *Alligator mississippiensis* K_Ca_5, and *Homo sapiens* K_Ca_2.2 were queried against MSAs generated from MMseqs2 (Steinegger and Söding, 2017) and structurally predicted 5 times with 6 recycles. Models ranked highest on ColabFold (highest ipTM score) were selected for downstream analysis. Contingent on available processing power, predicted structures were then relaxed using amber force fields (Eastman et al., 2017).

### Alignment of Transmembrane Segments (<u>Suppl. Files 1-8</u>)

All alignments were performed with the MAFFT (L-INS-i) web service in JalView. Initially, all species variants of a particular species are aligned to allow for extraction of each transmembrane segment based on the most evolutionarily similar structures. This allows for the alignments of transmembrane segments across all channels. A master alignment per transmembrane segment of each domain was done. From the master alignment, the specific alignments based on the groupings of voltage-gating, ion selectivity, and domain-swapping architecture were performed on the relevant TMS.

Sequence logos were made with WebLogo 3.1 (Crooks et al., 2004; Schneider and Stephens, 1990). The color coding is as follows: black - small, hydrophobic (GAPVILM); green - aromatic (FWY); orange - polar (STNQ); blue - positively charged (RKH); red - negatively charged (ED). Amino acids positions that were conserved in chemical and physical characteristics across channels in a particular grouping were highlighted for further investigation.

### ClinVar (<u>Suppl. Files 9-16</u>)

Clinically relevant missense mutations within the transmembrane segments for the 20 channels discussed in this paper were identified using ClinVar (Landrum et al., 2014). All human gene variants of each channel were used as search parameters (ie: KCNQ* for KCNQ channels, and KCNA* for Kv1-like channels). Those missense mutations annotated as benign, likely benign, likely pathogenic, and pathogenic were included in the database of this study. Mutated positions corresponding with highly conserved residues were highlighted with arrows (green for benign or likely benign; red for pathogenic or likely pathogenic).

## Author Contributions

Frank Yeh performed data compilation, sequence, structure, clinical variant analyses, wrote the manuscript, and managed the project. Arya Patel and Matthew Mun performed data compilation, sequence, structure, and clinical variant analyses, and edited the manuscript. Matthew Mun performed the AlphaFold2 structural predications. Youjin Oh performed sequence data compilation. Richard W. Aldrich wrote the manuscript.

