## Supplementary material for "Database Permeating through Time, Space, and Medicine: A sequence, structure, and clinical compilation and comparison of transmembrane amino acids in VGL ion channels": All referenced PDB files

### Supplemental Information

#### Table 1: FASTA Codes of All Sequences Used in Alignment

| **Species** | **KCNMA1** | **KCNT1** | **KCNU1** | **KCNN2** | **KCNN4** | **KCNA2** | **KCNB1** | **KCNC1** | **KCND1** |
| --- | --- | --- | --- | --- | --- | --- | --- | --- | --- |
| *Caspaspora owczarzaki* | No Return | [KJE92091.1](https://www-ncbi-nlm-nih-gov.ezproxy.lib.utexas.edu/protein/KJE92091.1?report=fasta) | No Return | No Return | No Return | No Return | [KJE94637.1](https://www-ncbi-nlm-nih-gov.ezproxy.lib.utexas.edu/protein/KJE94637.1?report=fasta) | No Return | No Return |
| *Salpingoeca rosetta* | No Return | [XP_004991549.1](https://www.ncbi.nlm.nih.gov/protein/XP_004991549.1?report=fasta) | No Return | No Return | No Return | No Return | [XP_004997201.1](https://www-ncbi-nlm-nih-gov.ezproxy.lib.utexas.edu/protein/XP_004997201.1?report=fasta) | No Return | No Return |
| *Bolinopsis microptera* | [XP_063691681.1](https://www.ncbi.nlm.nih.gov/protein/XP_063691681.1?report=genbank&log$=prottop&blast_rank=1&RID=CZ03N5XM016) | No Return | No Return | No Return | No Return | Provided by Jegla Lab (*Mnemiopsis leidyi*) | Provided by Jegla Lab (*Mnemiopsis leidyi*) | Provided by Jegla Lab (*Mnemiopsis leidyi*) | Provided by Jegla Lab (*Mnemiopsis leidyi*) |
| *Amphimedon queenslandica* | [XP_019857024.1](https://www-ncbi-nlm-nih-gov.ezproxy.lib.utexas.edu/protein/XP_019857024.1?report=fasta) | [XP_019852945.1](https://www-ncbi-nlm-nih-gov.ezproxy.lib.utexas.edu/protein/XP_019852945.1?report=fasta) | No Return | No Return | No Return | [XP_062513117.1 (*Corticium candelabrum*)](https://www.ncbi.nlm.nih.gov/protein/XP_062513117.1?report=fasta) | No Return | No Return | No Return |
| *Hydra vulgaris* | [XP_012564283.1](https://www-ncbi-nlm-nih-gov.ezproxy.lib.utexas.edu/protein/XP_012564283.1?report=fasta) | No Return | No Return | No Return | No Return | [XP_012556663](https://www-ncbi-nlm-nih-gov.ezproxy.lib.utexas.edu/protein/XP_012556663.1?report=fasta) | [XP_012559288.1](https://www-ncbi-nlm-nih-gov.ezproxy.lib.utexas.edu/protein/XP_012559288.1?report=fasta) | [XP_004208171.1](https://www.ncbi.nlm.nih.gov/protein/XP_004208171.1?report=fasta) | [XP_012556968.1](https://www.ncbi.nlm.nih.gov/protein/XP_012556968.1?report=fasta) |
| *Nematostella vectensis* | [XP_032219004.1 X1](https://www-ncbi-nlm-nih-gov.ezproxy.lib.utexas.edu/protein/XP_032219004.1?report=fasta) | [XP_032228039.1](https://www-ncbi-nlm-nih-gov.ezproxy.lib.utexas.edu/protein/XP_032228039.1?report=fasta) | No Return | [XP_032223175.1](https://www-ncbi-nlm-nih-gov.ezproxy.lib.utexas.edu/protein/XP_032223175.1?report=fasta) | No Return | [XP_001628953.2](https://www-ncbi-nlm-nih-gov.ezproxy.lib.utexas.edu/protein/XP_001628953.2?report=fasta) | [AJP09344.1](https://www-ncbi-nlm-nih-gov.ezproxy.lib.utexas.edu/protein/AJP09344.1?report=fasta) | [XP_001634787.2](https://www.ncbi.nlm.nih.gov/protein/XP_001634787.2?report=fasta) | [XP_032226945.1](https://www.ncbi.nlm.nih.gov/protein/XP_032226945.1?report=fasta) |
| *Helobdella robusta* | [XP_009020384.1](https://www.ncbi.nlm.nih.gov/protein/XP_009020384.1?report=fasta) | [XP_009016610.1](https://www-ncbi-nlm-nih-gov.ezproxy.lib.utexas.edu/protein/XP_009016610.1?report=fasta) | No Return | [XP_009011661.1](https://www-ncbi-nlm-nih-gov.ezproxy.lib.utexas.edu/protein/XP_009011661.1?report=fasta) | No Return | [XP_009032005.1](https://www.ncbi.nlm.nih.gov/protein/XP_009032005.1?report=fasta) | [XP_009029304.1](https://www-ncbi-nlm-nih-gov.ezproxy.lib.utexas.edu/protein/XP_009029304.1?report=fasta) | [XP_009015487.1](https://www.ncbi.nlm.nih.gov/protein/XP_009015487.1?report=fasta) | [XP_009011184.1](https://www.ncbi.nlm.nih.gov/protein/XP_009011184.1?report=fasta) |
| *Lottia gigantean* | [XP_009048035.1](https://www-ncbi-nlm-nih-gov.ezproxy.lib.utexas.edu/protein/XP_009048035.1?report=fasta) | [XP_009050341.1](https://www-ncbi-nlm-nih-gov.ezproxy.lib.utexas.edu/protein/XP_009050341.1?report=fasta) | No Return | [XP_009047871.1](https://www-ncbi-nlm-nih-gov.ezproxy.lib.utexas.edu/protein/XP_009047871.1?report=fasta) | No Return | [XP_009066018.1](https://www-ncbi-nlm-nih-gov.ezproxy.lib.utexas.edu/protein/XP_009066018.1?report=fasta) | [XP_009043764.1](https://www-ncbi-nlm-nih-gov.ezproxy.lib.utexas.edu/protein/XP_009043764.1?report=fasta) | [XP_009053105.1](https://www.ncbi.nlm.nih.gov/protein/XP_009053105.1?report=fasta) | [XP_009058094.1](https://www.ncbi.nlm.nih.gov/protein/XP_009058094.1?report=fasta) |
| *Aplysia californica* | [XP_035826261.1](https://www-ncbi-nlm-nih-gov.ezproxy.lib.utexas.edu/protein/XP_035826261.1?report=fasta) | [XP_012937020.1](https://www-ncbi-nlm-nih-gov.ezproxy.lib.utexas.edu/protein/XP_012937020.1?report=fasta) | No Return | [XP_035829626.1](https://www-ncbi-nlm-nih-gov.ezproxy.lib.utexas.edu/protein/XP_035829626.1?report=fasta) | No Return | [NP_001191634.1](https://www-ncbi-nlm-nih-gov.ezproxy.lib.utexas.edu/protein/NP_001191634.1?report=fasta) | [XP_005093895.1](https://www-ncbi-nlm-nih-gov.ezproxy.lib.utexas.edu/protein/XP_005093895.1?report=fasta) | [NP_001191546.1](https://www-ncbi-nlm-nih-gov.ezproxy.lib.utexas.edu/protein/NP_001191546.1?report=fasta) | [XP_005091742.1](https://www.ncbi.nlm.nih.gov/protein/XP_005091742.1?report=fasta) |
| *Schistosoma mansoni* | [XP_018647290.1](https://www-ncbi-nlm-nih-gov.ezproxy.lib.utexas.edu/protein/XP_018647290.1?report=fasta) | [PAA60613.1](https://www.ncbi.nlm.nih.gov/protein/PAA60613.1?report=fasta) | No Return | [XP_018649289.1](https://www-ncbi-nlm-nih-gov.ezproxy.lib.utexas.edu/protein/XP_018649289.1?report=fasta) | No Return | [XP_018651902.1](https://www-ncbi-nlm-nih-gov.ezproxy.lib.utexas.edu/protein/XP_018651902.1?report=fasta) | [XP_018646495.1](https://www-ncbi-nlm-nih-gov.ezproxy.lib.utexas.edu/protein/XP_018646495.1?report=fasta) | [KAK4467920.1 (*Schistosoma mekongi*)](https://www.ncbi.nlm.nih.gov/protein/KAK4467920.1?report=fasta) | [XP_018648931.1](https://www.ncbi.nlm.nih.gov/protein/XP_018648931.1?report=fasta) |
| *C. elegans* | [NP_001024259.1](https://www-ncbi-nlm-nih-gov.ezproxy.lib.utexas.edu/protein/NP_001024259.1?report=fasta) | [NP_001257146.1](https://www-ncbi-nlm-nih-gov.ezproxy.lib.utexas.edu/protein/NP_001257146.1?report=fasta) | No Return | [NP_001293203.1](https://www-ncbi-nlm-nih-gov.ezproxy.lib.utexas.edu/protein/NP_001293203.1?report=fasta) | No Return | [NP_871934.1](https://www.ncbi.nlm.nih.gov/protein/NP_871934.1?report=fasta) | [NP_001367760.1](https://www-ncbi-nlm-nih-gov.ezproxy.lib.utexas.edu/protein/NP_001367760.1?report=fasta) | [NP_001022089.1](https://www.ncbi.nlm.nih.gov/protein/NP_001022089.1?report=fasta) | [NP_500975.2](https://www.ncbi.nlm.nih.gov/protein/NP_500975.2?report=fasta) |
| *Drosophila melanogaster* | [NP_001262925.1](https://www-ncbi-nlm-nih-gov.ezproxy.lib.utexas.edu/protein/NP_001262925.1?report=fasta) | [NP_001097259.2](https://www-ncbi-nlm-nih-gov.ezproxy.lib.utexas.edu/protein/NP_001097259.2?report=fasta) | No Return | [NP_001284887.1](https://www-ncbi-nlm-nih-gov.ezproxy.lib.utexas.edu/protein/NP_001284887.1?report=fasta) | No Return | [NP_523393.3](https://www.ncbi.nlm.nih.gov/protein/NP_523393.3?report=fasta) | [NP_001189037.1](https://www-ncbi-nlm-nih-gov.ezproxy.lib.utexas.edu/protein/NP_001189037.1?report=fasta) | [NP_001137782.1](https://www.ncbi.nlm.nih.gov/protein/NP_001137782.1?report=fasta) | [NP_001097646.1](https://www.ncbi.nlm.nih.gov/protein/NP_001097646.1?report=fasta) |

Table 1 (*cont.*)

| **Species** | **KCNMA1** | **KCNT1** | **KCNU1** | **KCNN2** | **KCNN4** | **KCNA2** | **KCNB1** | **KCNC1** | **KCND1** |
| --- | --- | --- | --- | --- | --- | --- | --- | --- | --- |
| *Priapulus caudatus* | No Return | [XP_014668588.1](https://www-ncbi-nlm-nih-gov.ezproxy.lib.utexas.edu/protein/XP_014668588.1?report=fasta) | No Return | [XP_014663916.1](https://www-ncbi-nlm-nih-gov.ezproxy.lib.utexas.edu/protein/XP_014663916.1?report=fasta) | No Return | [XP_014681274.1](https://www-ncbi-nlm-nih-gov.ezproxy.lib.utexas.edu/protein/XP_014681274.1?report=fasta) | [XP_014670979.1](https://www-ncbi-nlm-nih-gov.ezproxy.lib.utexas.edu/protein/XP_014670979.1?report=fasta) | [XP_014674212.1](https://www.ncbi.nlm.nih.gov/protein/XP_014674212.1?report=fasta) | [XP_014670979.1](https://www-ncbi-nlm-nih-gov.ezproxy.lib.utexas.edu/protein/XP_014670979.1?report=fasta) |
| *S. purpuratus* | [XP_030835947.1](https://www-ncbi-nlm-nih-gov.ezproxy.lib.utexas.edu/protein/XP_030835947.1?report=fasta) | [XP_030853581.1](https://www-ncbi-nlm-nih-gov.ezproxy.lib.utexas.edu/protein/XP_030853581.1?report=fasta) | No Return | [XP_030832948.1](https://www-ncbi-nlm-nih-gov.ezproxy.lib.utexas.edu/protein/XP_030832948.1?report=fasta) | No Return | [XP_011672256.1](https://www.ncbi.nlm.nih.gov/protein/XP_011672256.1?report=fasta) | [XP_038057522.1 *(Patiria miniata*)](https://www.ncbi.nlm.nih.gov/protein/XP_038057522.1?report=fasta) | [XP_030845384.1](https://www.ncbi.nlm.nih.gov/protein/XP_030845384.1?report=fasta) | [XP_030856314.1](https://www.ncbi.nlm.nih.gov/protein/XP_030856314.1?report=fasta) |
| *Ciona intestinalis* | [XP_026693508.1](https://www-ncbi-nlm-nih-gov.ezproxy.lib.utexas.edu/protein/XP_026693508.1?report=fasta) | [XP_026694653.1](https://www-ncbi-nlm-nih-gov.ezproxy.lib.utexas.edu/protein/XP_026694653.1?report=fasta) | No Return | [XP_018671843.1](https://www-ncbi-nlm-nih-gov.ezproxy.lib.utexas.edu/protein/XP_018671843.1?report=fasta) | No Return | [XP_026692621.1](https://www-ncbi-nlm-nih-gov.ezproxy.lib.utexas.edu/protein/XP_026692621.1?report=fasta) | [XP_018671683.1](https://www-ncbi-nlm-nih-gov.ezproxy.lib.utexas.edu/protein/XP_018671683.1?report=fasta) | No Return | [XP_018671683.1](https://www-ncbi-nlm-nih-gov.ezproxy.lib.utexas.edu/protein/XP_018671683.1?report=fasta) |
| *B. floridae* | [XP_035669715.1](https://www-ncbi-nlm-nih-gov.ezproxy.lib.utexas.edu/protein/XP_035669715.1?report=fasta) | [XP_035684961.1](https://www-ncbi-nlm-nih-gov.ezproxy.lib.utexas.edu/protein/XP_035684961.1?report=fasta) | No Return | [XP_035667930.1](https://www-ncbi-nlm-nih-gov.ezproxy.lib.utexas.edu/protein/XP_035667930.1?report=fasta) | No Return | [XP_035672097.1](https://www-ncbi-nlm-nih-gov.ezproxy.lib.utexas.edu/protein/XP_035672097.1?report=fasta) | [XP_035666721.1](https://www-ncbi-nlm-nih-gov.ezproxy.lib.utexas.edu/protein/XP_035666721.1?report=fasta) | [XP_066275417.1 (*Branchiostoma laneolatum*)](https://www.ncbi.nlm.nih.gov/protein/XP_066275417.1?report=fasta) | [XP_035666721.1](https://www-ncbi-nlm-nih-gov.ezproxy.lib.utexas.edu/protein/XP_035666721.1?report=fasta) |
| *Petromyzon marinus* | [XP_032814990.1](https://www-ncbi-nlm-nih-gov.ezproxy.lib.utexas.edu/protein/XP_032814990.1?report=fasta) | [XP_032828853.1](https://www-ncbi-nlm-nih-gov.ezproxy.lib.utexas.edu/protein/XP_032828853.1?report=fasta) | No Return | [XP_032805199.1](https://www-ncbi-nlm-nih-gov.ezproxy.lib.utexas.edu/protein/XP_032805199.1?report=fasta) | No Return | [XP_032821220.1](https://www-ncbi-nlm-nih-gov.ezproxy.lib.utexas.edu/protein/XP_032821220.1?report=fasta) | [XP_032810624.1](https://www-ncbi-nlm-nih-gov.ezproxy.lib.utexas.edu/protein/XP_032810624.1?report=fasta) | [XP_032814254.1](https://www.ncbi.nlm.nih.gov/protein/XP_032814254.1?report=fasta) | [XP_032810624.1](https://www-ncbi-nlm-nih-gov.ezproxy.lib.utexas.edu/protein/XP_032810624.1?report=fasta) |
| *C. milii* | [XP_007896678.1](https://www-ncbi-nlm-nih-gov.ezproxy.lib.utexas.edu/protein/XP_007896678.1?report=fasta) | [XP_007898501.1](https://www-ncbi-nlm-nih-gov.ezproxy.lib.utexas.edu/protein/XP_007898501.1?report=fasta) | No Return | [XP_007895576.1](https://www-ncbi-nlm-nih-gov.ezproxy.lib.utexas.edu/protein/XP_007895576.1?report=fasta) | No Return | [NP_001280068.1](https://www-ncbi-nlm-nih-gov.ezproxy.lib.utexas.edu/protein/NP_001280068.1?report=fasta) | [XP_007885165.1](https://www-ncbi-nlm-nih-gov.ezproxy.lib.utexas.edu/protein/XP_007885165.1?report=fasta) | [XP_007885588.1](https://www.ncbi.nlm.nih.gov/protein/XP_007885588.1?report=fasta) | [XP_007885165.1](https://www-ncbi-nlm-nih-gov.ezproxy.lib.utexas.edu/protein/XP_007885165.1?report=fasta) |
| *Pogona vitticeps* | [XP_020651797.1](https://www.ncbi.nlm.nih.gov/protein/XP_020651797.1?report=fasta) | [XP_020664849.1](https://www-ncbi-nlm-nih-gov.ezproxy.lib.utexas.edu/protein/XP_020664849.1?report=fasta) | [XP_020641777.1](https://www-ncbi-nlm-nih-gov.ezproxy.lib.utexas.edu/protein/XP_020641777.1?report=fasta) | [XP_020635686.1](https://www-ncbi-nlm-nih-gov.ezproxy.lib.utexas.edu/protein/XP_020635686.1?report=fasta) | [XP_020663145.1](https://www-ncbi-nlm-nih-gov.ezproxy.lib.utexas.edu/protein/XP_020663145.1?report=fasta) | [XP_020665658.1](https://www-ncbi-nlm-nih-gov.ezproxy.lib.utexas.edu/protein/XP_020665658.1?report=fasta) | [XP_020668371.1](https://www-ncbi-nlm-nih-gov.ezproxy.lib.utexas.edu/protein/XP_020668371.1?report=fasta) | [XP_020666999.1](https://www.ncbi.nlm.nih.gov/protein/XP_020666999.1?report=fasta) | [XP_020668371.1](https://www-ncbi-nlm-nih-gov.ezproxy.lib.utexas.edu/protein/XP_020668371.1?report=fasta) |
| *Alligator m.* | [XP_019352722.1](https://www.ncbi.nlm.nih.gov/protein/XP_019352722.1?report=fasta) | [XP_019331175.1](https://www-ncbi-nlm-nih-gov.ezproxy.lib.utexas.edu/protein/XP_019331175.1?report=fasta) | [XP_019347112.1](https://www-ncbi-nlm-nih-gov.ezproxy.lib.utexas.edu/protein/XP_019347112.1?report=fasta) | [XP_006266937.3](https://www-ncbi-nlm-nih-gov.ezproxy.lib.utexas.edu/protein/XP_006266937.3?report=fasta) | [XP_019334954.1](https://www-ncbi-nlm-nih-gov.ezproxy.lib.utexas.edu/protein/XP_019334954.1?report=fasta) | [XP_019348081.1](https://www-ncbi-nlm-nih-gov.ezproxy.lib.utexas.edu/protein/XP_019348081.1?report=fasta) | [XP_019354264.1](https://www-ncbi-nlm-nih-gov.ezproxy.lib.utexas.edu/protein/XP_019354264.1?report=fasta) | [XP_019332660.1](https://www.ncbi.nlm.nih.gov/protein/XP_019332660.1?report=fasta) | [XP_019354264.1](https://www-ncbi-nlm-nih-gov.ezproxy.lib.utexas.edu/protein/XP_019354264.1?report=fasta) |
| *Gallus gallus* | [XP_015143651.1](https://www.ncbi.nlm.nih.gov/protein/XP_015143651.1?report=fasta) | [XP_015134798.1](https://www-ncbi-nlm-nih-gov.ezproxy.lib.utexas.edu/protein/XP_015134798.1?report=fasta) | [XP_040393979.1 (Cygnus olor)](https://www.ncbi.nlm.nih.gov/protein/XP_040393979.1?report=genbank&log$=protalign&blast_rank=1&RID=F3595JZJ016) | [XP_015136073.2](https://www-ncbi-nlm-nih-gov.ezproxy.lib.utexas.edu/protein/XP_015136073.2?report=fasta3) | No Return | [NP_989794.1](https://www-ncbi-nlm-nih-gov.ezproxy.lib.utexas.edu/protein/NP_989794.1?report=fasta) | [XP_015152169.1](https://www-ncbi-nlm-nih-gov.ezproxy.lib.utexas.edu/protein/XP_015152169.1?report=fasta) | [NP_004967.1](https://www-ncbi-nlm-nih-gov.ezproxy.lib.utexas.edu/protein/XP_004941482.1?report=fasta) | [XP_015152169.1](https://www-ncbi-nlm-nih-gov.ezproxy.lib.utexas.edu/protein/XP_015152169.1?report=fasta) |
| *Mus musculus* | [NP_034740.2](https://www-ncbi-nlm-nih-gov.ezproxy.lib.utexas.edu/protein/NP_034740.2?report=fasta) | [XP_006497940.2](https://www-ncbi-nlm-nih-gov.ezproxy.lib.utexas.edu/protein/XP_006497940.2?report=fasta) | [NP_032458.3](https://www-ncbi-nlm-nih-gov.ezproxy.lib.utexas.edu/protein/NP_032458.3?report=fasta) | [XP_030106191.1](https://www-ncbi-nlm-nih-gov.ezproxy.lib.utexas.edu/protein/XP_030106191.1?report=fasta) | [NP_001156982.1](https://www-ncbi-nlm-nih-gov.ezproxy.lib.utexas.edu/protein/NP_001156982.1?report=fasta) | [NP_032443.3](https://www-ncbi-nlm-nih-gov.ezproxy.lib.utexas.edu/protein/NP_032443.3?report=fasta) | [NP_001091998.1](https://www-ncbi-nlm-nih-gov.ezproxy.lib.utexas.edu/protein/NP_001091998.1?report=fasta) | [XP_006540709.1](https://www.ncbi.nlm.nih.gov/protein/XP_006540709.1?report=fasta) | [NP_001091998.1](https://www-ncbi-nlm-nih-gov.ezproxy.lib.utexas.edu/protein/NP_001091998.1?report=fasta) |
| *Homo sapiens* | [XP_016871702.1](https://www-ncbi-nlm-nih-gov.ezproxy.lib.utexas.edu/protein/XP_016871702.1?report=fasta) | [NP_065873.2](https://www.ncbi.nlm.nih.gov/protein/NP_065873.2?report=fasta) | [NP_001027006.2](https://www.ncbi.nlm.nih.gov/protein/NP_001027006.2?report=fasta) | [NP_001359162.1](https://www.ncbi.nlm.nih.gov/protein/NP_001359162.1?report=fasta) | [NP_002241.1](https://www.ncbi.nlm.nih.gov/protein/NP_002241.1?report=fasta) | [NP_004965.1](https://www.ncbi.nlm.nih.gov/protein/NP_004965.1?report=fasta) | [NP_004966.1](https://www.ncbi.nlm.nih.gov/protein/NP_004966.1?report=fasta) | [NP_004967.1](https://www.ncbi.nlm.nih.gov/protein/NP_004967.1?report=fasta) | [XP_024308146.1](https://www.ncbi.nlm.nih.gov/protein/XP_024308146.1?report=fasta) |

Table 1 (*cont.*)

| **Species** | **KCNQ** | **KCNH1, 5** | **KCNH2, 6, 7** | **KCNH3, 4, 8** | **HCN** | **CNGA** |
| --- | --- | --- | --- | --- | --- | --- |
| *Caspaspora owczarzaki* | No Return | No Return | No Return | No Return | No Return | No Return |
| *Salpingoeca rosetta* | No Return | No Return | No Return | No Return | [XP_004991538.1](https://www-ncbi-nlm-nih-gov.ezproxy.lib.utexas.edu/protein/XP_004991538.1?report=fasta) | [XP_004992545.1](https://www-ncbi-nlm-nih-gov.ezproxy.lib.utexas.edu/protein/XP_004992545.1?report=fasta) |
| *Bolinopsis microptera* | No Return | Provided by Jegla Lab (*Mnemiopsis leidyi*) | Provided by Jegla Lab (*Mnemiopsis leidyi*) | Provided by Jegla Lab (*Mnemiopsis leidyi*) | [XP_063686149.1](https://www.ncbi.nlm.nih.gov/protein/XP_063686149.1?report=genbank&log$=prottop&blast_rank=1&RID=CZ8WTUSA016) | [XP_063683221.1](https://www.ncbi.nlm.nih.gov/protein/XP_063683221.1?report=fasta) |
| *Amphimedon queenslandica* | No Return | [XP_065829990.1 (*Oscarella lobularis*)](https://www.ncbi.nlm.nih.gov/protein/XP_065829990.1?report=fasta) | [XP_062522508.1 (*Corticium candelabrum*)](https://www.ncbi.nlm.nih.gov/protein/XP_062522508.1?report=fasta) | No Return | [XP_065836457.1 (*Oscarella lobularis*)](https://www.ncbi.nlm.nih.gov/protein/XP_065836457.1?report=fasta) | [XP_003385808.1](https://www-ncbi-nlm-nih-gov.ezproxy.lib.utexas.edu/protein/XP_003385808.1?report=fasta) |
| *Hydra vulgaris* | [XP_012559369.1](https://www.ncbi.nlm.nih.gov/protein/XP_012559369.1?report=fasta) | [XP_012554704.1](https://www.ncbi.nlm.nih.gov/protein/XP_012554704.1?report=fasta) | [XP_002157210.3](https://www-ncbi-nlm-nih-gov.ezproxy.lib.utexas.edu/protein/XP_012559677.1?report=fasta) | [XP_057304855.1 (*Hydractinia symbiolongicarpus*)](https://www.ncbi.nlm.nih.gov/protein/XP_057304855.1?report=fasta) | [XP_012559677.1](https://www-ncbi-nlm-nih-gov.ezproxy.lib.utexas.edu/protein/XP_012559677.1?report=fasta) | [XP_012555740.1](https://www-ncbi-nlm-nih-gov.ezproxy.lib.utexas.edu/protein/XP_012555740.1?report=fasta) |
| *Nematostella vectensis* | [XP_001634862.2](https://www.ncbi.nlm.nih.gov/protein/XP_001634862.2?report=fasta) | [XP_032232877.1](https://www.ncbi.nlm.nih.gov/protein/XP_032232877.1?report=fasta) | [AHX24685.1](https://www-ncbi-nlm-nih-gov.ezproxy.lib.utexas.edu/protein/XP_001626416.2?report=fasta) | [XP_032218302.1](https://www.ncbi.nlm.nih.gov/protein/XP_032218302.1?report=fasta) | [XP_001626416.2](https://www-ncbi-nlm-nih-gov.ezproxy.lib.utexas.edu/protein/XP_001626416.2?report=fasta) | [XP_032231125.1](https://www-ncbi-nlm-nih-gov.ezproxy.lib.utexas.edu/protein/XP_032231125.1?report=fasta) |
| *Helobdella robusta* | [XP_009014289.1](https://www.ncbi.nlm.nih.gov/protein/XP_009014289.1?report=fasta) | [XP_009019489.1](https://www.ncbi.nlm.nih.gov/protein/XP_009019489.1?report=fasta) | [XP_009012155.1](https://www-ncbi-nlm-nih-gov.ezproxy.lib.utexas.edu/protein/XP_009021284.1?report=fasta) | [XP_009013566.1](https://www.ncbi.nlm.nih.gov/protein/XP_009013566.1?report=fasta) | [XP_009011849.1](https://www-ncbi-nlm-nih-gov.ezproxy.lib.utexas.edu/protein/XP_009011849.1?report=fasta) | [XP_009021284.1](https://www-ncbi-nlm-nih-gov.ezproxy.lib.utexas.edu/protein/XP_009021284.1?report=fasta) |
| *Lottia gigantean* | [XP_009054448.1](https://www.ncbi.nlm.nih.gov/protein/XP_009054448.1?report=fasta) | [XP_009063678.1](https://www.ncbi.nlm.nih.gov/protein/XP_009063678.1?report=fasta) | [XP_009061269.1](https://www.ncbi.nlm.nih.gov/protein/XP_012936081.1?report=fasta) | [XP_009048131.1](https://www.ncbi.nlm.nih.gov/protein/XP_009048131.1?report=fasta) | [XP_009054816.1](https://www-ncbi-nlm-nih-gov.ezproxy.lib.utexas.edu/protein/XP_009054816.1?report=fasta) | [XP_009047376.1](https://www-ncbi-nlm-nih-gov.ezproxy.lib.utexas.edu/protein/XP_009047376.1?report=fasta) |
| *Aplysia californica* | [XP_012934690.1](https://www.ncbi.nlm.nih.gov/protein/XP_012934690.1?report=fasta) | [XP_009063678.1](https://www.ncbi.nlm.nih.gov/protein/XP_009063678.1?report=fasta) | [XP_005107164.1](https://www-ncbi-nlm-nih-gov.ezproxy.lib.utexas.edu/protein/XP_018647037.1?report=fasta) | [XP_012936081.1](https://www.ncbi.nlm.nih.gov/protein/XP_012936081.1?report=fasta) | [NP_001191636.1](https://www-ncbi-nlm-nih-gov.ezproxy.lib.utexas.edu/protein/NP_001191636.1?report=fasta) | [XP_005093850.1](https://www-ncbi-nlm-nih-gov.ezproxy.lib.utexas.edu/protein/XP_005093850.1?report=fasta) |
| *Schistosoma mansoni* | [XP_018649462.1](https://www.ncbi.nlm.nih.gov/protein/XP_018649462.1?report=fasta) | [XP_018649958.1](https://www.ncbi.nlm.nih.gov/protein/XP_018649958.1?report=fasta) | [XP_018655373.1](https://www-ncbi-nlm-nih-gov.ezproxy.lib.utexas.edu/protein/NP_001033948.1?report=fasta) | [XP_018649331.1](https://www.ncbi.nlm.nih.gov/protein/XP_018649331.1?report=fasta) | [XP_018647037.1](https://www-ncbi-nlm-nih-gov.ezproxy.lib.utexas.edu/protein/XP_018647037.1?report=fasta) | [XP_018650675.1](https://www-ncbi-nlm-nih-gov.ezproxy.lib.utexas.edu/protein/XP_018650675.1?report=fasta) |
| *C. elegans* | [NP_001254403.1](https://www.ncbi.nlm.nih.gov/protein/NP_001254403.1?report=fasta) | [NP_001368173.1](https://www.ncbi.nlm.nih.gov/protein/NP_001368173.1?report=fasta) | [NP_503402.3](https://www.ncbi.nlm.nih.gov/protein/XP_030830580.1?report=fasta) | No Return | No Return | [5H3O_A](https://www-ncbi-nlm-nih-gov.ezproxy.lib.utexas.edu/protein/5H3O_A?report=fasta) |
| *Drosophila melanogaster* | [NP_001137631.1](https://www.ncbi.nlm.nih.gov/protein/NP_001137631.1?report=fasta) | [NP_001036275.1](https://www.ncbi.nlm.nih.gov/protein/NP_001036275.1?report=fasta) | [NP_001286814.1](https://www-ncbi-nlm-nih-gov.ezproxy.lib.utexas.edu/protein/XP_784539.3?report=fasta) | [NP_477009.1](https://www.ncbi.nlm.nih.gov/protein/NP_477009.1?report=fasta) | [NP_001033948.1](https://www-ncbi-nlm-nih-gov.ezproxy.lib.utexas.edu/protein/NP_001033948.1?report=fasta) | [NP_001188960.1](https://www-ncbi-nlm-nih-gov.ezproxy.lib.utexas.edu/protein/NP_001188960.1?report=fasta) |

Table 1 (*cont.*)

| **Species** | **KCNQ** | **KCNH1, 5** | **KCNH2, 6, 7** | **KCNH3, 4, 8** | **HCN** | **CNGA** |
| --- | --- | --- | --- | --- | --- | --- |
| *Priapulus caudatus* | [XP_014663789.1,](https://www.ncbi.nlm.nih.gov/protein/XP_014663789.1?report=fasta) | [XP_014677839.1](https://www.ncbi.nlm.nih.gov/protein/XP_014677839.1?report=fasta) | No Return | No Return | [XP_014679893.1](https://www-ncbi-nlm-nih-gov.ezproxy.lib.utexas.edu/protein/XP_014679893.1?report=fasta) | [XP_014676958.1](https://www-ncbi-nlm-nih-gov.ezproxy.lib.utexas.edu/protein/XP_014676958.1?report=fasta) |
| *S. purpuratus* | [XP_030845618.1](https://www.ncbi.nlm.nih.gov/protein/XP_030845618.1?report=fasta) | [XP_030855353.1](https://www.ncbi.nlm.nih.gov/protein/XP_030855353.1?report=fasta) | [XP_030852568.1](https://www-ncbi-nlm-nih-gov.ezproxy.lib.utexas.edu/protein/XP_035681348.1?report=fasta) | [XP_030830580.1](https://www.ncbi.nlm.nih.gov/protein/XP_030830580.1?report=fasta) | [NP_999729.1](https://www-ncbi-nlm-nih-gov.ezproxy.lib.utexas.edu/protein/NP_999729.1?report=fasta) | [XP_784539.3](https://www-ncbi-nlm-nih-gov.ezproxy.lib.utexas.edu/protein/XP_784539.3?report=fasta) |
| *Ciona intestinalis* | [NP_001153537.1](https://www.ncbi.nlm.nih.gov/protein/NP_001153537.1?report=fasta) | [XP_026694540.1](https://www.ncbi.nlm.nih.gov/protein/XP_026694540.1?report=fasta) | [XP_026692366.1](https://www.ncbi.nlm.nih.gov/protein/XP_007895827.1?report=fasta) | [XP_009859607.1](https://www.ncbi.nlm.nih.gov/protein/XP_009859607.1?report=fasta) | [XP_004226550.2](https://www-ncbi-nlm-nih-gov.ezproxy.lib.utexas.edu/protein/XP_004226550.2?report=fasta) | [XP_002124361.1](https://www-ncbi-nlm-nih-gov.ezproxy.lib.utexas.edu/protein/XP_002124361.1?report=fasta) |
| *B. floridae* | [XP_035663435.1](https://www.ncbi.nlm.nih.gov/protein/XP_035663435.1?report=fasta) | [XP_035695207.1](https://www.ncbi.nlm.nih.gov/protein/XP_035661277.1?report=fasta) | [XP_035688589.1](https://www-ncbi-nlm-nih-gov.ezproxy.lib.utexas.edu/protein/XP_020659343.1?report=fasta) | [XP_035661277.1](https://www.ncbi.nlm.nih.gov/protein/XP_035661277.1?report=fasta) | [XP_035667373.1](https://www-ncbi-nlm-nih-gov.ezproxy.lib.utexas.edu/protein/XP_035667373.1?report=fasta) | [XP_035681348.1](https://www-ncbi-nlm-nih-gov.ezproxy.lib.utexas.edu/protein/XP_035681348.1?report=fasta) |
| *Petromyzon marinus* | [XP_032804878.1](https://www.ncbi.nlm.nih.gov/protein/XP_032804878.1?report=fasta) | [XP_032820082.1](https://www-ncbi-nlm-nih-gov.ezproxy.lib.utexas.edu/protein/XP_032806347.1?report=fasta) | [XP_032801658.1](https://www-ncbi-nlm-nih-gov.ezproxy.lib.utexas.edu/protein/XP_006272154.1?report=fasta) | [XP_032818464.1](https://www.ncbi.nlm.nih.gov/protein/XP_032818464.1?report=fasta) | [XP_032806347.1](https://www-ncbi-nlm-nih-gov.ezproxy.lib.utexas.edu/protein/XP_032806347.1?report=fasta) | [XP_032836778.1](https://www-ncbi-nlm-nih-gov.ezproxy.lib.utexas.edu/protein/XP_032836778.1?report=fasta) |
| *C. milii* | [XP_007892746.1](https://www.ncbi.nlm.nih.gov/protein/XP_007892746.1?report=fasta) | [XP_007902076.1](https://www-ncbi-nlm-nih-gov.ezproxy.lib.utexas.edu/protein/XP_007889307.1?report=fasta) | [XP_007888033.1](https://www.ncbi.nlm.nih.gov/protein/NP_038597.2?report=fasta) | [XP_007895827.1](https://www.ncbi.nlm.nih.gov/protein/XP_007895827.1?report=fasta) | [XP_007908262.1](https://www-ncbi-nlm-nih-gov.ezproxy.lib.utexas.edu/protein/XP_007908262.1?report=fasta) | [XP_007889307.1](https://www-ncbi-nlm-nih-gov.ezproxy.lib.utexas.edu/protein/XP_007889307.1?report=fasta) |
| *Pogona vitticeps* | [XP_020644480.1](https://www.ncbi.nlm.nih.gov/protein/XP_020644480.1?report=fasta) | [XP_020665464.1](https://www.ncbi.nlm.nih.gov/protein/XP_019351032.1?report=fasta) | [XP_020655898.1](https://www.ncbi.nlm.nih.gov/protein/XP_065829990.1?report=fasta) | [XP_020662997.1](https://www.ncbi.nlm.nih.gov/protein/XP_020662997.1?report=fasta) | [XP_020659343.1](https://www-ncbi-nlm-nih-gov.ezproxy.lib.utexas.edu/protein/XP_020659343.1?report=fasta) | [XP_020665211.1](https://www-ncbi-nlm-nih-gov.ezproxy.lib.utexas.edu/protein/XP_020665211.1?report=fasta) |
| *Alligator m.* | [XP_014457848.2](https://www.ncbi.nlm.nih.gov/protein/XP_014457848.2?report=fasta) | [XP_014454245.1](https://www.ncbi.nlm.nih.gov/protein/XP_040520526.1?report=fasta) | [XP_019351032.1](https://www.ncbi.nlm.nih.gov/protein/XP_009063678.1?report=fasta) | [XP_019353285.1](https://www.ncbi.nlm.nih.gov/protein/XP_019353285.1?report=fasta) | [XP_014459956.1](https://www-ncbi-nlm-nih-gov.ezproxy.lib.utexas.edu/protein/XP_014459956.1?report=fasta) | [XP_006272154.1](https://www-ncbi-nlm-nih-gov.ezproxy.lib.utexas.edu/protein/XP_006272154.1?report=fasta) |
| *Gallus gallus* | [XP_421022.5](https://www.ncbi.nlm.nih.gov/protein/XP_421022.5?report=fasta) | [XP_040529709.1](https://www.ncbi.nlm.nih.gov/protein/XP_036016335.1?report=fasta) | [XP_015136636.3](https://www.ncbi.nlm.nih.gov/protein/XP_014677839.1?report=fasta) | [XP_040520526.1](https://www.ncbi.nlm.nih.gov/protein/XP_040520526.1?report=fasta) | [XP_040536079.1](https://www-ncbi-nlm-nih-gov.ezproxy.lib.utexas.edu/protein/XP_040536079.1?report=fasta) | [NP_990552.1](https://www-ncbi-nlm-nih-gov.ezproxy.lib.utexas.edu/protein/NP_990552.1?report=fasta) |
| *Mus musculus* | [XP_006508551.2](https://www.ncbi.nlm.nih.gov/protein/XP_006508551.2?report=fasta) | [NP_766393.2](https://www.ncbi.nlm.nih.gov/protein/XP_032232877.1?report=fasta) | [NP_038597.2](https://www-ncbi-nlm-nih-gov.ezproxy.lib.utexas.edu/protein/XP_032806347.1?report=fasta) | [XP_036016335.1](https://www.ncbi.nlm.nih.gov/protein/XP_036016335.1?report=fasta) | [NP_001074661.1](https://www-ncbi-nlm-nih-gov.ezproxy.lib.utexas.edu/protein/NP_001074661.1?report=fasta) | [NP_001268939.1](https://www-ncbi-nlm-nih-gov.ezproxy.lib.utexas.edu/protein/NP_001268939.1?report=fasta) |
| *Homo sapiens* | [NP_000209.2](https://www.ncbi.nlm.nih.gov/protein/NP_000209.2?report=fasta) | [NP_647479.2](https://www.ncbi.nlm.nih.gov/protein/NP_647479.2?report=fasta) | [NP_000229.1](https://www.ncbi.nlm.nih.gov/protein/NP_000229.1?report=fasta) | [NP_653234.2](https://www.ncbi.nlm.nih.gov/protein/NP_653234.2?report=fasta) | [NP_005468.1](https://www.ncbi.nlm.nih.gov/protein/NP_005468.1?report=fasta) | [NP_005131.1](https://www.ncbi.nlm.nih.gov/protein/NP_005131.1?report=fasta) |

Table 1 (*cont.*)

| **Species** | **Nav1** | **Cav1** | **Cav2** | **Cav3** | **NALCN** |
| --- | --- | --- | --- | --- | --- |
| *Caspaspora owczarzaki* | No Return | No Return | No Return | No Return | No Return |
| *Salpingoeca rosetta* | [XP_004998888.1](https://www.ncbi.nlm.nih.gov/protein/XP_004998888.1?report=genbank&log$=prottop&blast_rank=1&RID=M297SFSC013) | [UIX25862.1](https://www.ncbi.nlm.nih.gov/protein/UIX25862.1?report=genbank&log$=prottop&blast_rank=1&RID=39J1DN2Y013) | XP_004989719.1 | [XP_004995501.1](https://www.ncbi.nlm.nih.gov/protein/XP_004995501.1?report=fasta) | No Return |
| *Bolinopsis microptera* | [XP_063675154.1](https://www.ncbi.nlm.nih.gov/protein/XP_063675154.1?report=genbank&log$=prottop&blast_rank=1&RID=CZC34JF5016) | [QOY24596.1](https://www.ncbi.nlm.nih.gov/protein/QOY24596.1?report=fasta) | XP_063686312.1 | No Return | No Return |
| *Amphimedon queenslandica* | No Return | [XP_019855146.1](https://www.ncbi.nlm.nih.gov/protein/XP_019855146.1?report=genbank&log$=prottop&blast_rank=1&RID=39GJMBN6016) | XP_019855140.1 | No Return | [XP_019854422.1](https://www.ncbi.nlm.nih.gov/protein/XP_019854422.1?report=genbank&log$=prottop&blast_rank=1&RID=39NNVK4B016) |
| *Hydra vulgaris* | [XP_012554828.1](https://www.ncbi.nlm.nih.gov/protein/XP_012554828.1?report=genbank&log$=prottop&blast_rank=1&RID=M0208PBN013) | [XP_012567147.1](https://www.ncbi.nlm.nih.gov/protein/XP_012567147.1?report=genpept) | XP_047138459.1 | [XP_065658360.1](https://www.ncbi.nlm.nih.gov/protein/XP_065658360.1?report=fasta) | [XP_012565236.1](https://www.ncbi.nlm.nih.gov/protein/XP_012565236.1?report=genbank&log$=prottop&blast_rank=1&RID=39NCHUYF013) |
| *Nematostella vectensis* | [AEX00070.1](https://www.ncbi.nlm.nih.gov/protein/AEX00070.1?report=genbank&log$=prottop&blast_rank=1&RID=M02WX7U601R) | [XP_032219777.1](https://www.ncbi.nlm.nih.gov/protein/XP_032219777.1?report=genpept) | XP_048587791.1 | [XP_032231233.2](https://www.ncbi.nlm.nih.gov/protein/XP_032231233.2?report=fasta) | [XP_001637238.2](https://www.ncbi.nlm.nih.gov/protein/XP_001637238.2?report=genbank&log$=prottop&blast_rank=1&RID=39NHF4MU013) |
| *Helobdella robusta* | [XP_009012996.1](https://www.ncbi.nlm.nih.gov/protein/XP_009012996.1?report=genbank&log$=prottop&blast_rank=1&RID=M28NF986013) | [XP_009029798.1](https://www.ncbi.nlm.nih.gov/protein/XP_009029798.1?report=genbank&log$=prottop&blast_rank=1&RID=39GRBFM5013) | XP_009012628.1 | [XP_009014808.1](https://www.ncbi.nlm.nih.gov/protein/XP_009014808.1?report=fasta) | [XP_009025299.1](https://www.ncbi.nlm.nih.gov/protein/XP_009025299.1?report=genbank&log$=prottop&blast_rank=1&RID=39NTZX6C013) |
| *Lottia gigantean* | [XP_009063823.1](https://www.ncbi.nlm.nih.gov/protein/XP_009063823.1?report=genbank&log$=prottop&blast_rank=1&RID=M2S6U5F9016) | [XP_009057793.1](https://www.ncbi.nlm.nih.gov/protein/XP_009057793.1?report=genbank&log$=prottop&blast_rank=1&RID=39KH3A7C016) | XP_009057793.1 | [XP_009061843.1](https://www.ncbi.nlm.nih.gov/protein/XP_009061843.1?report=fasta) | [XP_009049022.1](https://www.ncbi.nlm.nih.gov/protein/XP_009049022.1?report=genbank&log$=prottop&blast_rank=1&RID=4HKUZB0H013) |
| *Aplysia californica* | [NP_001191637.1](https://www.ncbi.nlm.nih.gov/protein/NP_001191637.1?report=genbank&log$=prottop&blast_rank=1&RID=KZW4WUCB013) | [XP_035829232.1](https://www.ncbi.nlm.nih.gov/protein/XP_035829232.1?report=genbank&log$=prottop&blast_rank=1&RID=39EBVBV8016) | AVD53847.1 | [XP_035826040.1](https://www.ncbi.nlm.nih.gov/protein/XP_035826040.1?report=fasta) | [XP_012934773.1](https://www.ncbi.nlm.nih.gov/protein/XP_012934773.1?report=genbank&log$=prottop&blast_rank=1&RID=39N147G8016) |
| *Schistosoma mansoni* |  | [AAK84312.1](https://www.ncbi.nlm.nih.gov/protein/AAK84312.1?report=genbank&log$=prottop&blast_rank=1&RID=39KBG5UF013) | AAK84311.1 | No Return | [XP_018651519.1](https://www.ncbi.nlm.nih.gov/protein/XP_018651519.1?report=genbank&log$=prottop&blast_rank=1&RID=4HKN3M1V013) |
| *C. elegans* |  | [NP_741442.1](https://www.ncbi.nlm.nih.gov/protein/NP_741442.1?report=genbank&log$=prottop&blast_rank=1&RID=39FK3585016) | No Return | [NP_001367691.1](https://www.ncbi.nlm.nih.gov/protein/NP_001367691.1?report=fasta) | [NP_741413.2](https://www.ncbi.nlm.nih.gov/protein/NP_741413.2?report=genbank&log$=prottop&blast_rank=1&RID=39N7DBC7013) |
| *Drosophila melanogaster* | [NP_001188614.1](https://www-ncbi-nlm-nih-gov.ezproxy.lib.utexas.edu/protein/NP_001188614.1?report=fasta) | [NP_001260480.2](https://www.ncbi.nlm.nih.gov/protein/NP_001260480.2?report=fasta) | NP_001014732.1 | [NP_001259267.1](https://www.ncbi.nlm.nih.gov/protein/NP_001259267.1?report=fasta) | [NP_727772.3](https://www.ncbi.nlm.nih.gov/protein/NP_727772.3?report=fasta) |

Table 1 (*cont.*)

| **Species** | **Nav1** | **Cav1** | **Cav2** | **Cav3** | **NALCN** |
| --- | --- | --- | --- | --- | --- |
| *Priapulus caudatus* | [XP_014665068.1](https://www.ncbi.nlm.nih.gov/protein/XP_014665068.1?report=genbank&log$=prottop&blast_rank=1&RID=M2RVCTHZ013) | [XP_014661450.1](https://www.ncbi.nlm.nih.gov/protein/XP_014661450.1?report=genbank&log$=prottop&blast_rank=1&RID=39K61242016) | XP_014668456.1 | [XP_014676246.1](https://www.ncbi.nlm.nih.gov/protein/XP_014676246.1?report=fasta) | [XP_014674294.1](https://www.ncbi.nlm.nih.gov/protein/XP_014674294.1?report=genbank&log$=prottop&blast_rank=1&RID=4HKE7V11016) |
| *S. purpuratus* | [XP_030833314.1](https://www.ncbi.nlm.nih.gov/protein/XP_030833314.1?report=genbank&log$=prottop&blast_rank=1&RID=KZSMM8BG01R) | [XP_030827885.1](https://www.ncbi.nlm.nih.gov/protein/XP_030827885.1?report=genbank&log$=prottop&blast_rank=1&RID=39E3KMW9013) | XP_030853655.1 | [XP_030839905.1](https://www.ncbi.nlm.nih.gov/protein/XP_030839905.1?report=fasta) | [XP_030837794.1](https://www.ncbi.nlm.nih.gov/protein/XP_030837794.1?report=genbank&log$=prottop&blast_rank=1&RID=39MVB7K2016) |
| *Ciona intestinalis* | [XP_026691928.1](https://www.ncbi.nlm.nih.gov/protein/XP_026691928.1?report=genbank&log$=prottop&blast_rank=1&RID=M2A22Y12016) | [XP_026689962.1](https://www.ncbi.nlm.nih.gov/protein/XP_026689962.1?report=genbank&log$=prottop&blast_rank=1&RID=39JUJST4013) | XP_026693133.1 | [XP_018667817.1](https://www.ncbi.nlm.nih.gov/protein/XP_018667817.1?report=fasta) | [XP_009858187.2](https://www.ncbi.nlm.nih.gov/protein/XP_009858187.2?report=genbank&log$=prottop&blast_rank=1&RID=39PZ75ZK016) |
| *B. floridae* | [XP_035662890.1](https://www.ncbi.nlm.nih.gov/protein/XP_035662890.1?report=genbank&log$=prottop&blast_rank=1&RID=M2RUERBY013) | [XP_035676519.1](https://www.ncbi.nlm.nih.gov/protein/XP_035676519.1?report=genbank&log$=prottop&blast_rank=1&RID=39K1KBW3013) | XP_035684367.1 | [XP_035695929.1](https://www.ncbi.nlm.nih.gov/protein/XP_035695929.1?report=fasta) | [XP_035663373.1](https://www.ncbi.nlm.nih.gov/protein/XP_035663373.1?report=genbank&log$=prottop&blast_rank=1&RID=4HK2PNJS013) |
| *Petromyzon marinus* | [XP_032831283.1](https://www.ncbi.nlm.nih.gov/protein/XP_032831283.1?report=genbank&log$=prottop&blast_rank=1&RID=M29N5XFK013) | [XP_032822465.1](https://www.ncbi.nlm.nih.gov/protein/XP_032822465.1?report=genbank&log$=prottop&blast_rank=1&RID=39JKKR7B013) | XP_032829228.1 | [XP_032832080.1](https://www.ncbi.nlm.nih.gov/protein/XP_032832080.1?report=fasta) | [XP_032834381.1](https://www.ncbi.nlm.nih.gov/protein/XP_032834381.1?report=genbank&log$=prottop&blast_rank=1&RID=39PR1VCW013) |
| *C. milii* | [XP_007888018.2](https://www.ncbi.nlm.nih.gov/protein/XP_007888018.2?report=genbank&log$=prottop&blast_rank=1&RID=M29C6AMM016) | [XP_042197958.1](https://www.ncbi.nlm.nih.gov/protein/XP_042197958.1?report=genbank&log$=prottop&blast_rank=1&RID=39JDUV7X016) | XP_042200570.1 | [XP_042198287.1](https://www.ncbi.nlm.nih.gov/protein/XP_042198287.1?report=fasta) | [XP_007885224.1](https://www.ncbi.nlm.nih.gov/protein/XP_007885224.1?report=genbank&log$=prottop&blast_rank=1&RID=39PE6SMA013) |
| *Pogona vitticeps* | [XP_020645073.1](https://www.ncbi.nlm.nih.gov/protein/XP_020645073.1?report=genbank&log$=prottop&blast_rank=1&RID=M2TU67DK016) | [XP_020660962.1](https://www.ncbi.nlm.nih.gov/protein/XP_020660962.1?report=genbank&log$=prottop&blast_rank=1&RID=39KVR2KG013) | XP_020656028.1 | [XP_020670763.1](https://www.ncbi.nlm.nih.gov/protein/XP_020670763.1?report=fasta) | [XP_020651220.1](https://www.ncbi.nlm.nih.gov/protein/XP_020651220.1?report=genbank&log$=prottop&blast_rank=1&RID=4HM1U61H016) |
| *Alligator m.* | [XP_019335955.1](https://www.ncbi.nlm.nih.gov/protein/XP_019335955.1?report=genbank&log$=prottop&blast_rank=1&RID=M2TYXPEE013) | [XP_014464919.1](https://www.ncbi.nlm.nih.gov/protein/XP_014464919.1?report=genbank&log$=prottop&blast_rank=1&RID=39M44B3K013) | XP_059588283.1 | [XP_059588087.1](https://www.ncbi.nlm.nih.gov/protein/XP_059588087.1?report=fasta) | [XP_019342449.1](https://www.ncbi.nlm.nih.gov/protein/XP_019342449.1?report=genbank&log$=prottop&blast_rank=1&RID=4HM5WSAU013) |
| *Gallus gallus* | [XP_003641634.4](https://www.ncbi.nlm.nih.gov/protein/XP_003641634.4?report=genbank&log$=prottop&blast_rank=1&RID=M2U922E8016) | [NP_001384628.1](https://www.ncbi.nlm.nih.gov/protein/NP_001384628.1?report=genbank&log$=prottop&blast_rank=1&RID=39MEWHS0016) | XP_046784616.1 | [XP_040505818.1](https://www.ncbi.nlm.nih.gov/protein/XP_040505818.1?report=fasta) | [XP_015130472.1](https://www.ncbi.nlm.nih.gov/protein/XP_015130472.1?report=genbank&log$=prottop&blast_rank=1&RID=4HMBZRA7013) |
| *Mus musculus* | [NP_001300926.1](https://www.ncbi.nlm.nih.gov/protein/NP_001300926.1?report=genbank&log$=prottop&blast_rank=1&RID=M2SD2ASX016) | [NP_001074492.1](https://www.ncbi.nlm.nih.gov/protein/NP_001074492.1?report=genbank&log$=prottop&blast_rank=1&RID=39KMW0GM01N) | XP_006530659.1 | [XP_011246988.1](https://www.ncbi.nlm.nih.gov/protein/XP_011246988.1?report=fasta) | [XP_036014576.1](https://www.ncbi.nlm.nih.gov/protein/XP_036014576.1?report=genbank&log$=prottop&blast_rank=1&RID=4HKZ1XX6016) |
| *Homo sapiens* | [NP_001189364.1](https://www.ncbi.nlm.nih.gov/protein/NP_001189364.1?report=fasta) | [NP_000060.2](https://www.ncbi.nlm.nih.gov/protein/NP_000060.2?report=fasta) | NP_075461.2 | [NP_061496.2](https://www.ncbi.nlm.nih.gov/protein/NP_061496.2?report=fasta) | [XP_024305104.1](https://www.ncbi.nlm.nih.gov/protein/XP_024305104.1?report=fasta) |

#### Table 2: PDB Codes for each Channel Structure

|  | KCNMA1 | KCNT1 | KCNU1 | KCNN2 | KCNN4 | KCNA2 | KCNB1 | KCNC1 | KCND1 | KCNQ |
| --- | --- | --- | --- | --- | --- | --- | --- | --- | --- | --- |
| PDB IDs | [6V38](https://www.rcsb.org/structure/6V38) | [5U70](https://www.rcsb.org/structure/5U70) | AlphaFold2  Prediction | AlphaFold2  Prediction | [6CNO](https://www.rcsb.org/structure/6CNO) | [3LUT](https://www.rcsb.org/structure/3LUT) | [8SD3](https://www.rcsb.org/structure/8SD3) | [8QUD](https://www.rcsb.org/structure/8QUD) | [7UK5](https://www.rcsb.org/structure/7UK5) | [5VMS](https://www.rcsb.org/structure/5VMS) |

|  | KCNH1+5 | KCNH2,6,7 | KCNH3,4,8 | HCN | CNGA | Nav 1.1 | Cav 1.1 | Cav 2.1 | Cav 3.1 | NALCN |
| --- | --- | --- | --- | --- | --- | --- | --- | --- | --- | --- |
| PDB  IDs | [6PBX](https://www.rcsb.org/structure/6PBX) | [5VA2](https://www.rcsb.org/structure/5VA2) | AlphaFold2  Prediction | [5U6O](https://www.rcsb.org/structure/5U6O) (VSD)  [8T50](https://www.rcsb.org/structure/8T50)  (Pore) | [7LFY](https://www.rcsb.org/structure/7LFY) | [7DTD](https://www.rcsb.org/structure/7DTD) | [6JPA](https://www.rcsb.org/structure/6JPA) | [8X90](https://www.rcsb.org/structure/8X90) | [6KZO](https://www.rcsb.org/structure/6KZO) | [6XIW](https://www.rcsb.org/structure/6XIW) |
